# Crystal structure of a bacterial CNNM magnesium transporter

**DOI:** 10.1101/2021.02.11.430706

**Authors:** Yu Seby Chen, Guennadi Kozlov, Joshua Armitano, Brandon E. Moeller, Rayan Fakih, Ahmed Rohaim, Benoit Roux, John E. Burke, Kalle Gehring

## Abstract

CBS-pair domain divalent metal cation transport mediators (CNNMs) are a broadly conserved family of integral membrane proteins with close to 90,000 protein sequences known. CNNM proteins are associated with Mg^2+^ transport but it is not known if they mediate transport themselves or regulate other transporters. Here, we determined the crystal structure of an archaeal CNNM protein with Mg^2+^-ATP bound. The structure reveals a novel transmembrane fold for the DUF21 domain, the largest family of domains of unknown function. The protein has a negatively charged cavity that penetrates halfway through the membrane suggesting it functions as a cation transporter. The cytosolic portion of the protein is comprised of highly charged four-helix bundle and a CBS-pair domain. HDX-MS experiments, molecular dynamics, and additional crystal structures show that the cytosolic domains undergo large conformational changes upon nucleotide binding suggesting a mechanism of regulation shared between human and bacterial orthologs. The molecular characterization of CNNM proteins has profound implications for understanding their biological functions in human diseases, including cancer, and in animals, bacteria and plants.

## Introduction

CNNMs (<u>C</u>BS-pair domai<u>n</u> divalent catio<u>n</u> transport <u>m</u>ediators) are a conserved family of integral membrane proteins implicated in Mg^2+^ homeostasis and divalent cation transport (Funato and Miki, 2019; Wang et al., 2003). Also known as ancient conserved domain proteins (ACDPs), they are found in all species from bacteria, yeast, plants and animals (Fig. 1a). Although they have been genetically associated with Mg^2+^ transport, their biochemical function is disputed. Some researchers claim they are Mg^2+^ transporters while other believe they act as sensors (Arjona and de Baaij, 2018; Funato et al., 2018). In humans, mutations in CNNM proteins are linked to two genetic diseases: dominant *CNNM2* mutations cause hypomagnesemia (Arjona et al., 2014; Stuiver et al., 2011) while recessive mutations in *CNNM4* are associated with Jalili syndrome (Daneshmandpour et al., 2019). CNNMs are also implicated in cancer through binding of PRLs, highly oncogenic protein phosphatases involved in metastasis (Funato et al., 2014; Hardy et al., 2015). PRL binding inhibits CNNM-associated Mg^2+^ efflux and is in turn is regulated by magnesium levels (Gulerez et al., 2016; Hirata et al., 2014; Kozlov et al., 2020). In bacteria and yeast, mutations in CNNMs confer resistance to toxic metals and sensitivity to elevated levels of Mg^2+^.

**Figure 1.**
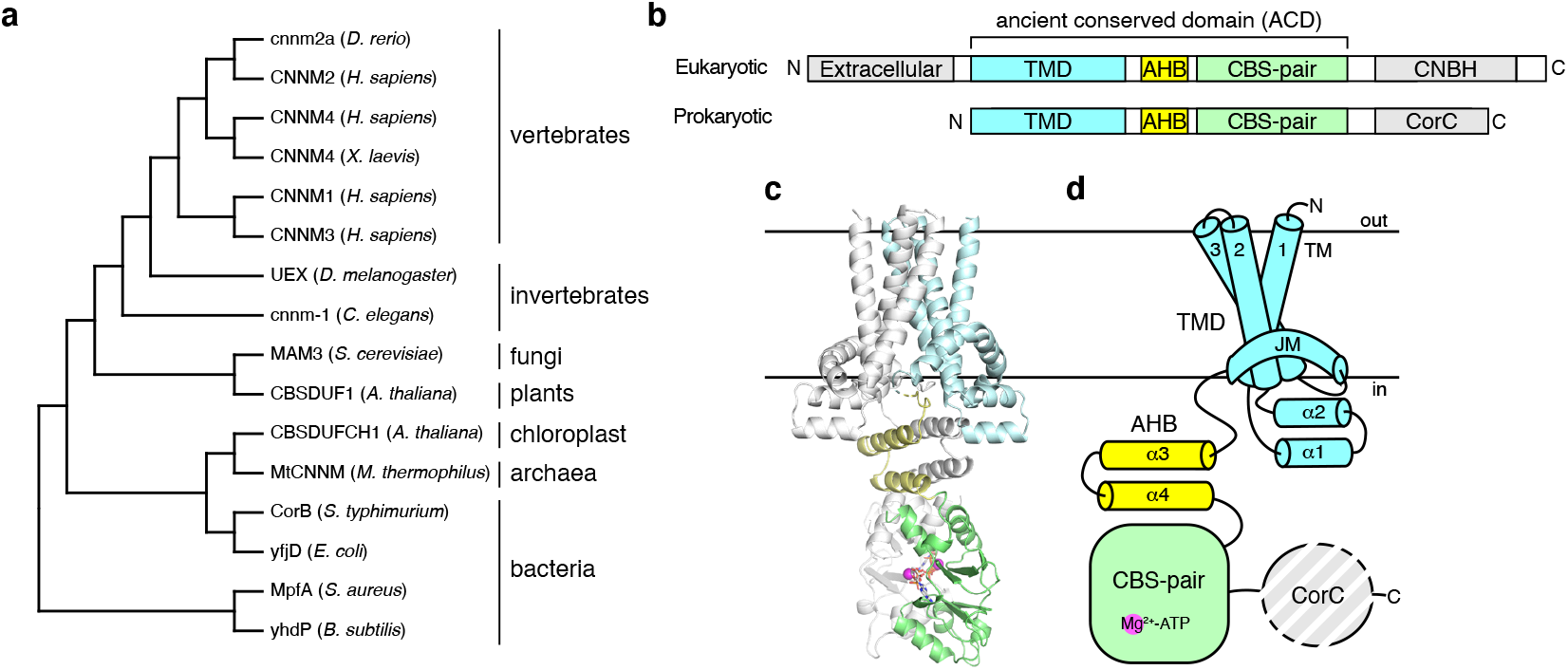
Crystal structure of MtCNNMΔC bound to Mg^2+^-ATP. **a**, Phylogenetic analysis of CNNM orthologs generated using neighbor-joining method. **b**, Domain organization of eukaryotic and prokaryotic CNNMs. TMD, transmembrane domain; AHB, acidic helical bundle; CNBH, cyclic nucleotide-binding homology domain; CorC, cobalt resistance C domain. **c**, Crystal structure of MtCNNM without the C-terminal CorC domain as a homodimer. One chain is colored by domains. **d**, Topology of a MtCNNM monomer showing the transmembrane and juxtamembrane helices of the TMD (residues 1-154, *cyan*), the two helices of the AHB (residues 166-199, *yellow*), the Mg^2+^- ATP-binding CBS-pair (residues 200-324, *green*), and the CorC domain (residues 325-426, *grey*).

Structurally, CNNMs are defined by a conserved core consisting of a transmembrane domain and a cytosolic cystathionine-β-synthase (CBS)-pair domain (Fig. 1b). The transmembrane domain constitutes the largest family of protein domains of unknown function (DUF21) in the Pfam database (El-Gebali et al., 2019). The Interpro database lists 87,000 known and predicted CNNM proteins (Blum et al., 2021). Outside of the core, eukaryotic and prokaryotic CNNMs diverge. Eukaryotes have an extracellular domain and cytosolic C-terminal cyclic-nucleotide binding homology (CNBH) domains. Prokaryotes lack the extracellular domain and have a distinct C-terminal domain, CorC.

CBS-pair domains are composed of tandem CBS motifs and found in a wide variety of proteins, including the bacterial Mg^2+^ transporter MgtE, where they mediate dimerization and Mg^2+^-ATP binding (Ereno-Orbea et al., 2013). The CBS-pair domains of CNNMs have been characterized structurally and shown to undergo large conformational changes upon Mg^2+^-ATP binding (Chen et al., 2020; Corral-Rodriguez et al., 2014). The domains are the site of PRL binding that regulates the activity of CNNM proteins in humans (Gulerez et al., 2016; Kozlov et al., 2020).

Here, we determined the crystal structure of an archaeal CNNM in the presence and absence of Mg^2+^-ATP. The structure reveals the protein acts to transport cations with a highly acidic cavity and multiple regulatory cation binding sites. The cytosolic domains undergo large conformational changes upon nucleotide binding suggesting a shared mechanism of regulation from human to bacterial orthologs. The demonstration that CNNMs are directly involved in Mg^2+^ transport is of primary importance for understanding their biological function in cancer and Mg^2+^ homeostasis.

## Results

### Domain architecture

We screened CNNM orthologs and identified the protein from the archaeon *Methanoculleus thermophilus* (MtCNNM) as the most promising candidate for structural studies (Fig. S1). The full-length protein did not yield suitable crystals but deletion of an internal loop and the C-terminal CorC domain (MtCNNMΔC_Δloop_) allowed determination of structure of the transmembrane and regulatory cytoplasmic domain (Fig. S2a, Table S1). MtCNNMΔC_Δloop_ forms a homodimer with each protomer consisting of three distinct regions: a transmembrane domain (TMD), an acidic helical bundle (AHB), and a CBS-pair domain (Fig. 1c-d). The TMD and AHB are unique with no similarity to known structures.

Although MtCNNM is present as a dimer, it does not show C2 symmetry (Fig. S3). The individual TMD, AHB and CBS-pair domains form symmetric dimers, but their arrangement is not. Overlaying the TMD of two polypeptide chains reveals asymmetry in the positions of the cytosolic domains, primarily due to differences in the linker between the TMD and AHB. This suggests the interface between the transmembrane and cytosolic domains is dynamic and may play a role in regulation of transporter activity.

### Structure reveals MCNNM is a cation transporter

The TMD is a dimer with each chain composed of three transmembrane helices (TM1-3), a pair of short helices exposed on the intracellular side, and a juxtamembrane helix (JM) that forms a belt-like structure (Fig. 2a). The TMD forms a large, negatively charged cavity in the membrane with a bound Na^+^ ion (Fig. 2b). Dimerization is predominantly formed by hydrophobic contacts of TM2 and TM3 of each protomer which also form the walls of the inward facing cavity (Fig. 2a). The cavity has a maximum diameter of approximately 10 Å and is lined with conserved polar and acidic residues (Fig. 2c). Additional negative charge is present on the outer surface. Well-defined electron density is observed in the cavity, which was identified as Na^+^ based on the interatomic distances and valance screening analysis (Zheng et al., 2014) (Fig. 2d). The Na^+^ is coordinated by hydroxyl groups of Ser21, Ser25, and Ser71; carboxyl group of Glu111; and the main-chain carbonyl groups of Ser21 and Gly110. Residues contributing to ion interactions are strongly conserved across species, suggesting these are conserved features of the channel (Fig. S4).

**Figure 2.**
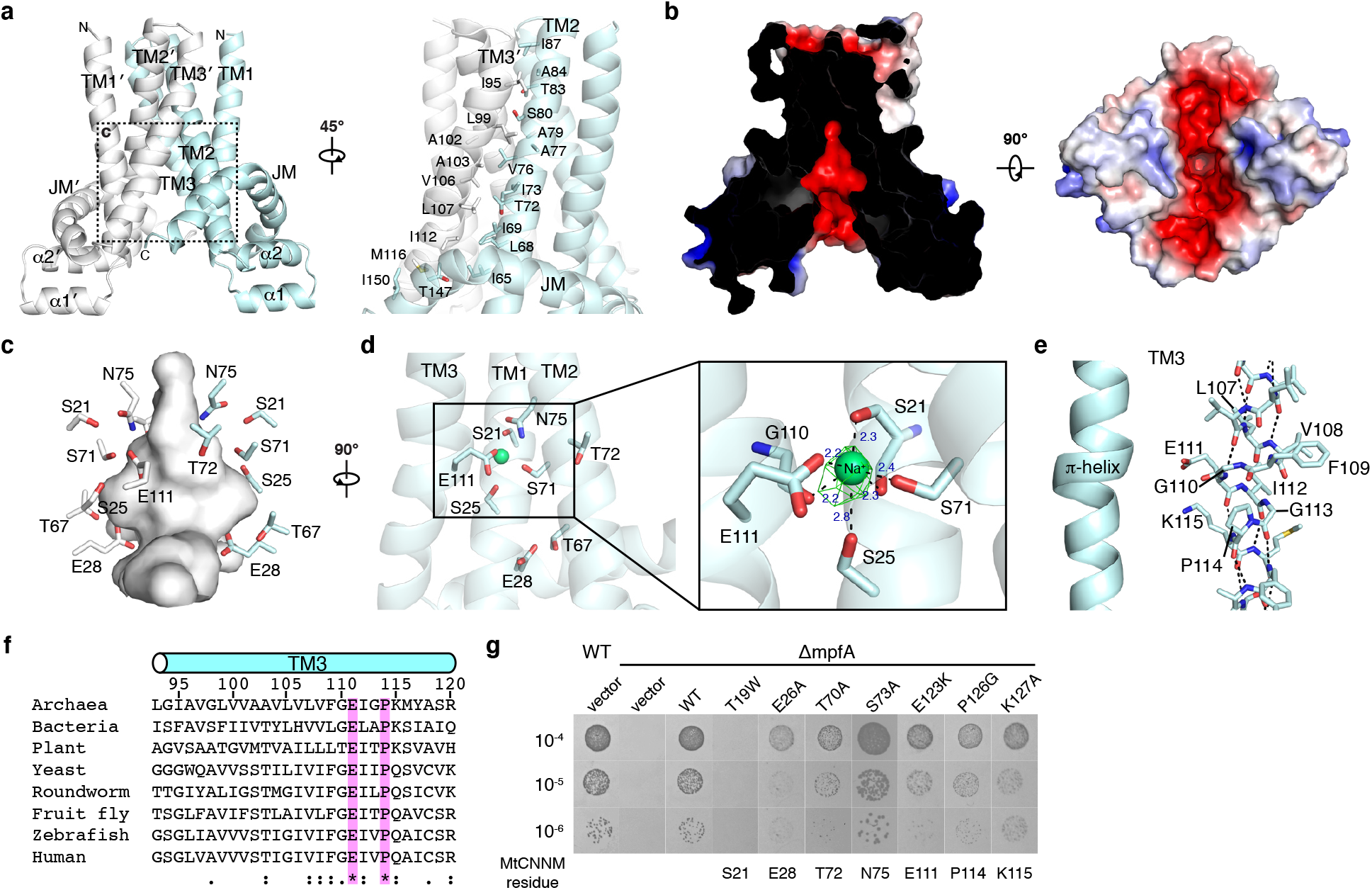
TM domain has negatively charged cavity on intracellular side. **a**, TM domain homodimerizes with interface formed by TM2 and TM3 of each protomer. **b**, Electrostatic surface potential representation (±5 kT e^-1^) of MtCNNM TM domain of cross-sectional (*left*) and intracellular (*right*) views of the acidic cavity. **c**, Close-up view of the residues forming the acidic cavity. **d**, A sodium ion (Na^+^) bound in the cavity with F_o_-F_c_ omit map contoured at 4.0 σ. **e**, π-helical turn preceding Pro114 in TM3. **f**, Conservation of residues in the π-helix turn from archaea to humans. **g**, Mutational analysis of MpfA, a CNNM ortholog in *S. aureus*. T19W, corresponding to mutation of S21 in MtCNNM, blocked complementation of a chromosomal deletion (ΔMpfA) and growth on plates with 140 mM MgCl_2_.

A second conserved feature is a π-helical turn in TM3 that forms a bulge in the helix and contributes to Na^+^ binding (Fig. 2e & S2a). The residues surrounding the helical turn are highly conserved with Glu111 and Pro114 completely invariant from archaea to humans (Fig. 2f). Glu111 points toward the negative cavity and is involved in coordination of the Na^+^ ion, whereas Pro114 acts as a helix-breaker to allow the shift in hydrogen-bonding in the π-helical turn.

We used an *in vivo* complementation assay with MpfA (SA0657), a CNNM ortholog in *Staphylococcus aureus*, to assess the functional importance of residues in the TMD. MpfA is 27% identical to MtCNNM and required for growth of *S. aureus* in the presence of high concentrations of Mg^2+^ (Armitano et al., 2016; Trachsel et al., 2019). In the complementation assay, growth of a ΔMpfA strain is rescued with a plasmid expressing wild-type MpfA (Fig. 2g & S5). Mutation of MpfA residues equivalent to Thr72, Ser75, Glu111, Pro114 and Lys115 of MtCNNM fully restored growth, while Glu28 partially restored growth, and Ser21 blocked rescue. This last mutation corresponds to the human CNNM2 mutation (S269W) identified in patients with hypomagnesemia (Arjona et al., 2014).

### Lipid binding interface

The re-entrant JM helix following TM3 wraps around TM1 and TM2 like a belt and interacts with hydrophobic residues in the protein and phospholipid headgroups in the membrane. The helix is rich in aromatic residues and amphipathic. We observed electron density for ten UDM molecules surrounding the JM helix (Fig. 3a-b). A helixturn-helix motif (α1 & α2) connects TM1 and TM2 and interacts with the JM helix through hydrophobic contacts. The solvent-exposed portions contain several basic residues, which may be involved in binding phospholipid headgroups. Supporting this, strong density was observed in this region and assigned as a sulfate molecule due to its presence in the crystallization buffer (Fig. S2b).

**Figure 3.**
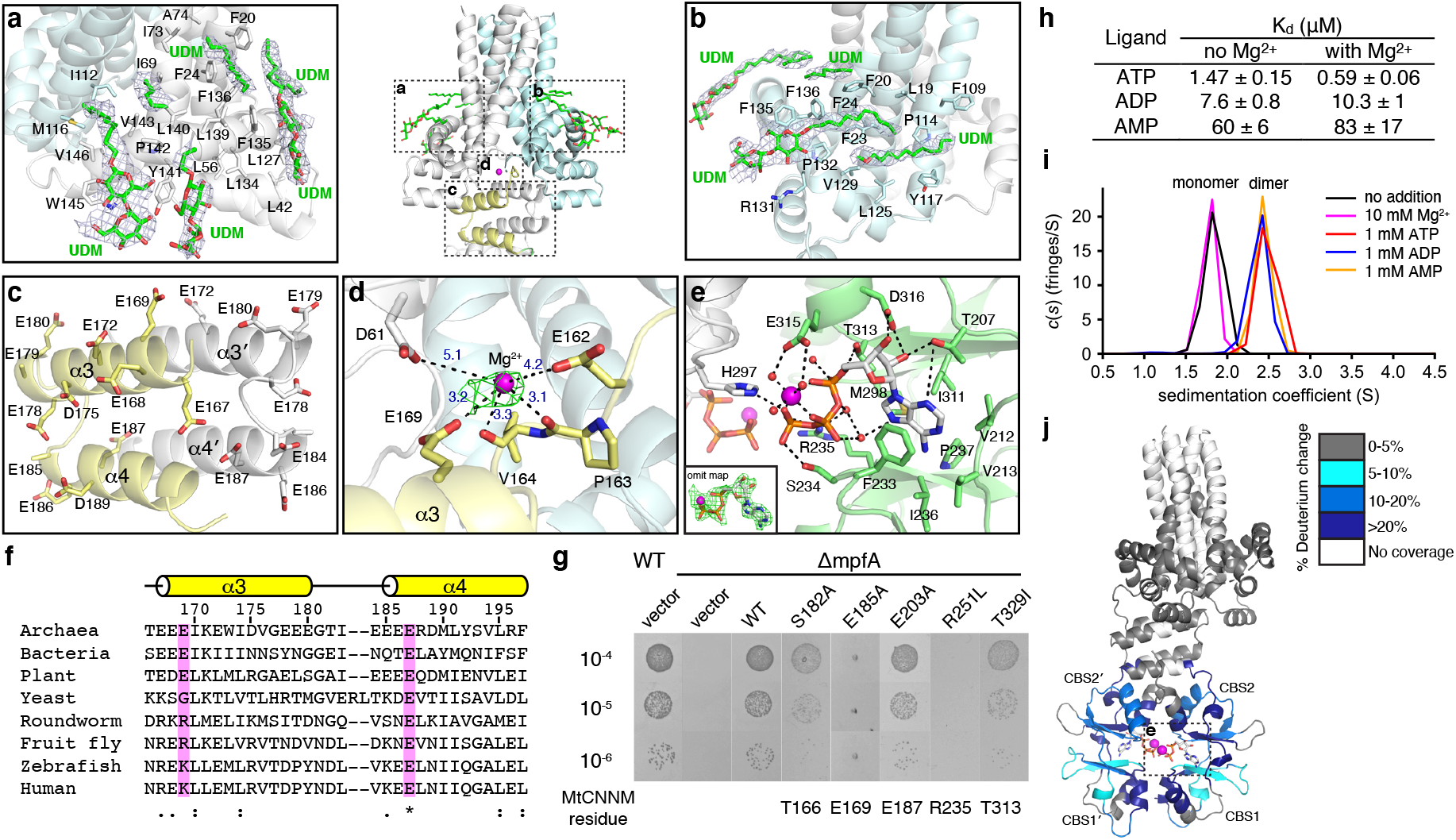
Juxtamembrane helix and conformational change in cytosolic domains. **a-b**, UDM detergent molecules bound by the juxtamembrane helix and 2F_o_-F_c_ map contoured at 1.0 σ. **c**, Conservation of acidic residues in the AHB domain. **d**, Hydrated magnesium bound between TMD and AHB and F_o_-F_c_ omit map contoured at 3 σ. **e**, Structural basis of Mg^2+^-ATP binding. Mg^2+^ ions and water molecules are shown in magenta and red, respectively. The Mg^2+^-ATP F_o_-F_c_ omit map was contoured at 3.0 σ. **f**, Sequence conservation of conserved glutamates in the AHB. **g**, Mutational analysis of MpfA in *S. aureus*. Mutation of a glutamate in the AHB and the CBS-pair domain arginine involved in ATP binding blocked growth. **h**, Affinities of MtCNNMΔC to adenosine nucleotides with and without 50 mM Mg^2+^ measured by ITC. **i**, Dimerization of the MtCNNM CBS-pair domain in the presence of 1 mM adenosine nucleotides as measured by SV-AUC experiments. **j**, Conformational change in the MtCNNMΔC in the presence and absence of Mg^2+^-ATP measured by HDX-MS. Regions that showed significant decreases in exchange (defined as >5%, 0.3 kDa, and a Student’s *t*-test p<0.01) in the presence of Mg^2+^-ATP are colored blue. Peptides in the CBS-pair domain dimerization interface show the most protection from deuterium exchange upon Mg^2+^-ATP binding.

### Acidic helical bundle

An unexpected feature of the MtCNNM structure is the existence of the four-helix AHB domain following the TMD (Fig. 3c). The linker between the juxtamembrane helix and AHB differs in the two protomers. It is extended and partially disordered in one and coiled in the other. The AHB itself is a symmetric dimer composed of two helices from each protomer and highly negatively charged. Thirteen of the 31 residues are acidic with three stretches of three or more consecutive glutamic acids. These likely bind Mg^2+^ and, indeed, we observed density around Glu169, that could be modelled as a hydrated Mg^2+^ ion (Fig. 3d). The bound Mg^2+^ is anchored by carboxylate groups of three acidic residues (Asp61, Glu162, and Glu169) and by the backbone carbonyl atoms of Pro163 and Val164. The Mg^2+^-O bond distances are between 3.1 Å and 5.1 Å, which is consistent with a hydrated Mg^2+^ ion.

The negative charge of the AHB is well conserved across evolution with the exception of yeast and roundworms (Fig. 3f). At least two invariant glutamate residues are present in all bacterial homologs, including the Mg^2+^-binding Glu169. To assess the importance of these, we turned again to the *S. aureus* complementation assay. Mutation of E185A in MpfA, which corresponds to loss of MtCNNM Glu169, was unable to support growth whereas mutations S182A and E203A in the MpfA AHB were able to complement the chromosomal MpfA deletion (Fig. 3g & S5).

### Structural basis of Mg^2+^-ATP binding by the CBS-pair domain

We crystallized the isolated MtCNNM CBS-pair to visualize the detailed interactions of Mg^2+^-ATP binding at higher resolution (Fig. 3e; Table S1). As observed in the structure with the TMD and AHB domains, two molecules of Mg^2+^-ATP are bound to the central cavity of the CBS-pair dimer. The adenine bases are sandwiched between Phe233 and Ile311 in a hydrophobic pocket comprising Met298, Phe237, Ile236, Val212, and Val213. The ATP ribose forms hydrogen bonds with the side chains of Thr207, Asp316, and Thr313. The phosphate groups are stabilized with Arg235 and Ser234 with the bound Mg^2+^ coordinated in an octahedral arrangement with three phosphates and three water molecules.

We tested the functional importance of these contacts in MpfA with a focus on mutations found in human diseases. Arg235 corresponds to the R407L mutation in CNNM4 that causes Jalili syndrome (Parveen et al., 2019) and Thr313 corresponds to T568I in CNNM2 that causes a dominant form of hypomagnesemia (Stuiver et al., 2011). Testing the mutations in MpfA demonstrated the importance of Arg235 but not Thr313 for rescue of the Mg^2+^-dependent growth defect (Fig. 3g & S5).

Measurements of the affinity of adenosine nucleotides by isothermal titration calorimetry (ITC) showed MtCNNMΔC has the highest affinity for ATP, followed by ADP and AMP (Fig. 3h & S6a). While addition of Mg^2+^ has a relatively large effect on binding of ATP to the isolated CBS-pair domain in human CNNMs (Chen et al., 2020; Hirata et al., 2014), the effect was more muted with MtCNNM. In presence of Mg^2+^, the affinity for ATP was increased three-fold with MtCNNMΔC and ten-fold with the isolated MtCNNM_CBS_ domain (Fig. S6b).

We used analytical ultracentrifugation (AUC) to characterize the effect of ATP binding on MtCNNM. As observed for human CNNMs, dimerization of the CBS-pair domain was tightly coupled with Mg^2+^-ATP binding. While the MtCNNM_CBS_ in the absence of ATP sedimented as a monomer, addition of adenine nucleotides triggered dimerization (Fig. 3i & S7). Addition of Mg^2+^ alone had no effect on the sedimentation of MtCNNM_CBS_.

We turned to hydrogen-deuterium exchange mass spectrometry (HDX-MS) to detect conformational changes in full-length MtCNNM upon ATP binding and deletion of the CorC domain. MtCNNM and MtCNNMΔC were reconstituted into nanodiscs and hydrogen exchange monitored in the presence and absence of Mg^2+^-ATP (Fig. S8a).

While peptide coverage of the hydrophobic TMD was limited, coverage of the cytosolic domains was excellent and revealed large conformational changes. Addition of Mg^2+^- ATP significantly reduced exchange in the CBS-pair domain dimerization interface and ATP-binding site (Fig. 3j & S8b-c). Similar changes were observed with both the fulllength and truncated protein, which is consistent with the absence of a role of C-terminal domain in Mg^2+^-ATP binding. This is different from human CNNMs where the C-terminal CNBH domain promotes CBS-pair domain dimerization and Mg^2+^-ATP binding. Comparison of full-length and truncated MtCNNM showed changes in the second CBS motif of the CBS-pair domain, suggesting that this is the primary site of interaction of the CorC domain.

### Conformational changes in MtCNNM upon loss of Mg^2+^-ATP binding

Further insight into the conformation in the absence of bound ATP came from the crystal structure of an MtCNNMΔC mutant that is unable to bind ATP (Fig. 4a, Table S1). The R235L mutant shows major rearrangements in the structure of the cytosolic domains with dissociation of the AHB and CBS-pair dimers and generation of a novel domain-domain contacts (Fig. 4b). In agreement with the AUC results, loss of ATP binding disrupted dimerization of the CBS-pair domain. Release of this interface allows the negatively charged helices of AHB to bind to the CBS-pair domain ATP-binding site, dramatically changing the interface between the TMD and cytosolic domains (Fig. 4c). The arrangement of AHB helices and CBS-pair domain is symmetric with about 10% of the solvent surface (~835 Å^2^) of each protomer buried in the interface. Dimerization is primarily driven by contacts between the AHB helices and CBS2 motif (Fig. 4d). The

**Figure 4.**
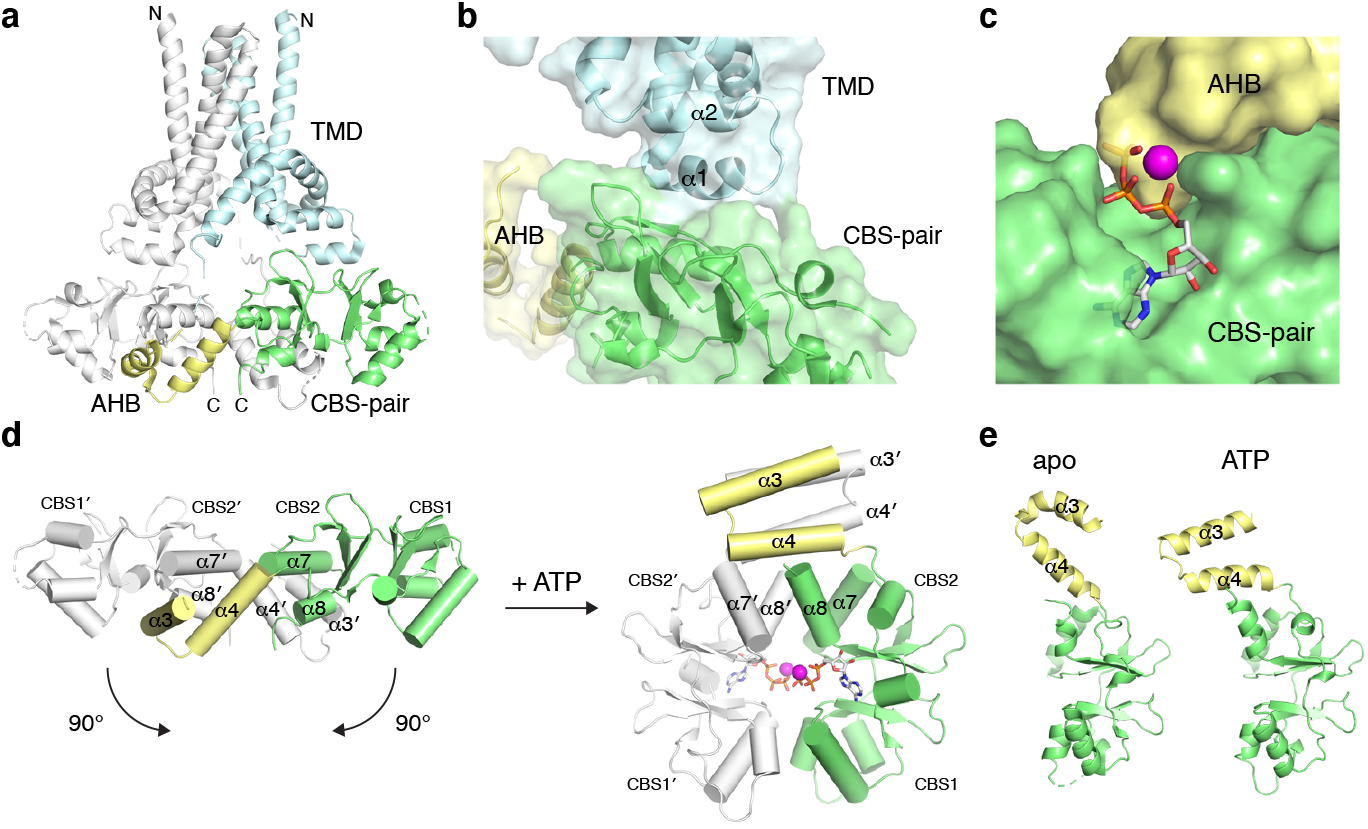
Apo conformation of MtCNNMΔC captured by R235L mutant. **a**, Overall structure of MtCNNMΔC R235L mutant. **b**, Contact surface between TMD and CBS-pair domain. **c**, AHB binds to CBS-pair domain and competes for Mg^2+^-ATP site. **d**, Large conformational change in the AHB and CBS-pair domains upon Mg^2+^-ATP binding. **e**, Structural comparison of apo and ATP-bound conformations.

CBS-pair domains from both structures are very similar with differences limited to the AHB helices (Fig. 4e). The R235L mutation is not directly involved in contacts between the domains and appears to act simply through the disruption of nucleotide binding.

Despite the large change in the cytosolic domains, the structure of the TMD was unaffected by the loss of Mg^2+^-ATP; the backbone atom RMSD between the two structures is 0.65 Å (Fig. 4). The negatively charged cavity is present in the R235L structure and fully accessible to ions on the cytosolic side. One consequence of the conformational change in the cytosolic domains is that the linkers following the TMD no longer come together to join the four-helical AHB. This might constrain the TMD from adopting an open conformation to allow ions access from the extracellular side. The ATP-less structure is more symmetric overall than the Mg^2+^-ATP structure but there are still differences between the protomers. In one, an exposed helix from the TMD binds the CBS2 motif while the other chain has no direct contact.

### Molecular modeling of MtCNNM

We used molecular modeling to improve our understanding of mutations in human (HsCNNM) proteins that cause diseases. Previous attempts to predict the structure of HsCNNM2 were confounded by the hydrophobicity of the HsCNNM2 JM helix (de Baaij et al., 2012) (Fig. S9a). Using the Mg^2+^-ATP bound structure of MtCNNM, we generated a homology model of HsCNNM2, which showed good geometry and allowed the positions of the helices in the TMD domains to be reliably positioned (Fig. 5a & S9). The main differences lie in the length of linkers connecting the TM helices. The HsCNNM2 model reproduced the π-helix in TM3, the Na^+^ ion binding site, and the presence buried polar residues (e.g. Thr331, Asp335, and Ser 348), supporting the importance of these features for CNNM function (Fig. 5b).

**Figure 5.**
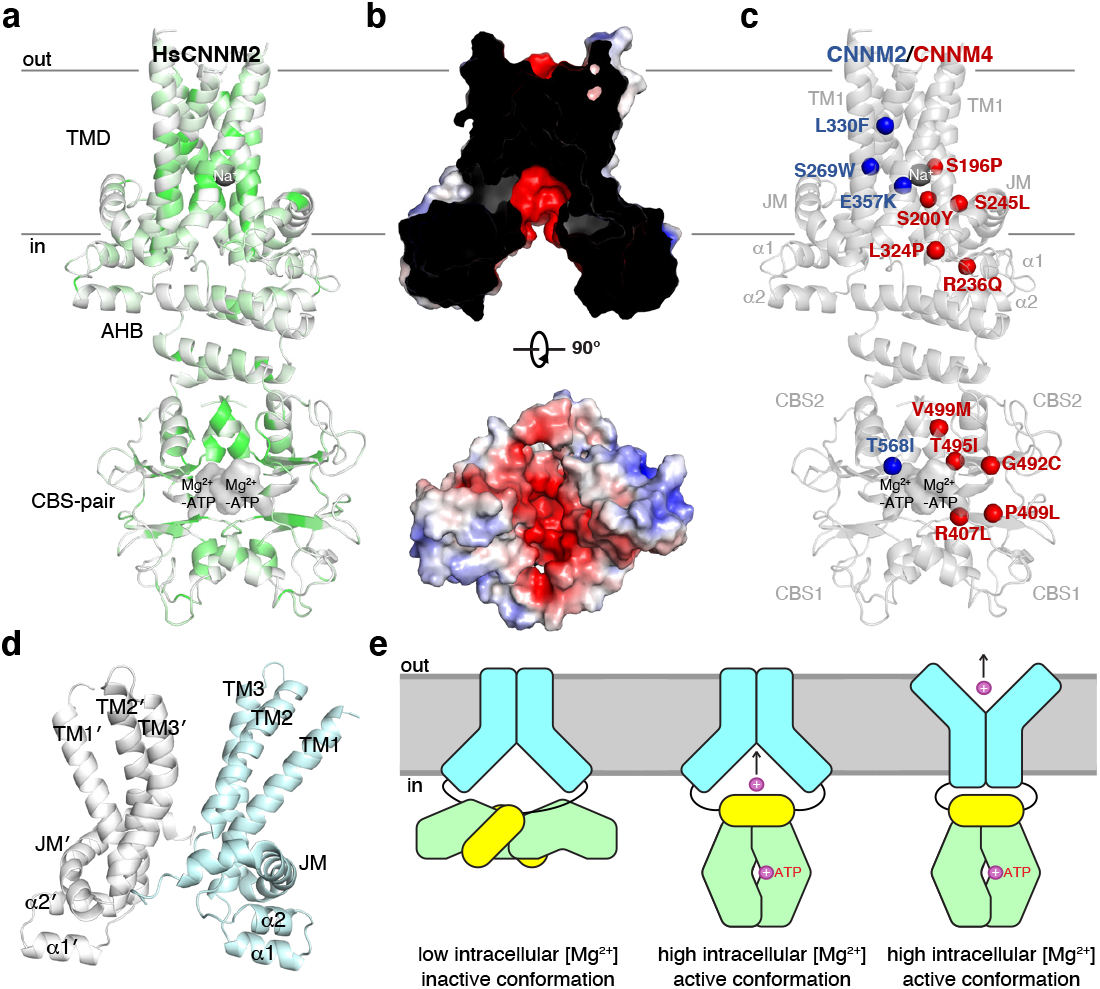
Molecular modeling. **a**, Homology model of human CNNM2 (HsCNNM2) color-coded according to identity to CNNMs shown in Fig. S4. The HsCNNM2 domains are 26% identical to MtCNNM. Conservation is highest around the Na^+^ binding site in the TMD, negatively charged cavity and the CBS-pair dimerization interface. **b**, Electrostatic surface potential representation (±5 kT e^-1^) of HsCNNM TMD in crosssectional (*top*) and intracellular (*bottom*) views. **c**, Mutations in CNNM2 and CNNM4 responsible for hypomagnesemia and Jalili syndrome cluster around the core of the TMD and nucleotide-binding site. For clarity, mutations are only shown on one chain. **d**, Outward-facing conformation generated by MD simulations. **e**, Proposed model of MtCNNM transport and regulation.

We used the model to understand the molecular basis for human disease mutations (Fig. 5c). Four missense mutations in HsCNNM2 are responsible for hypomagnesemia. T568I in the CBS-pair domain prevents ATP binding and Mg^2+^ transport (Chen et al., 2020; Hirata et al., 2014). In the TMD, S269W and L330F both likely disrupt TM helix packing and E357K is a charge reversal mutation located on the surface of the negatively charged cavity. While the MpfA mutant equivalent to E357K was active in *S. aureus*, the mutant corresponding to S269W was not (Fig. 2g). The high degree of sequence conservation (60% identity) between HsCNNM2 and HsCNNM4 also allowed us to use the model to interpret HsCNNM4 mutations that cause Jalili syndrome. Of particular interest are mutations that occur in the TMD. S196P and S200Y both map to the negatively charged cavity (Fig. 5c). HsCNNM4 S196P affects the same residue as CNNM2 S269W, confirming the importance of this position for function. R236Q, S245L, and L324P are located at the inner surface of the membrane, in TM3, and the juxtamembrane helix, respectively.

We next used molecular dynamics simulations to study asymmetry and conformational changes in the MtCNNM structure with Mg^2+^-ATP. As expected from the asymmetry in the structure, the interface between the TMD and cytosolic domains was flexible and large movements were observed (Video S1). At the end of the simulation, the conformations of the two protomers were essentially switched while the individual domains all retained their symmetry.

Targeted molecular dynamics was used to gain insight into the conformational changes in the TMD required for ion transport (Video S2). Pulling a Na^+^ through the TMD in 10 ns simulation generated a possible outward-facing conformation (Fig. 5d). The conserved residues Glu10, Ser80 and The83, which are buried in the membrane in the inward-facing conformation, become solvent exposed and available for ion binding. The conservation of the π-helix and buried polar residues suggest that the mechanism of ion transport is conserved across all CNNM family members.

## Discussion

Mg^2+^ is the most abundant divalent cation inside cells and essential for a wide variety of biochemical and enzymatic reactions (de Baaij et al., 2015). Despite this, our structural knowledge of Mg^2+^ ion transporter is limited to three proteins: MgtE, CorA, and TRPM7 (Duan et al., 2018; Eshaghi et al., 2006; Hattori et al., 2007; Lunin et al., 2006; Payandeh and Pai, 2006). The structure of MtCNNM defines a new class of Mg^2+^ transporter, highly conserved, and present in all organisms.

CNNMs had previously been suggested to be too small function in ion transport due to the small number of predicted TM helices (Arjona and de Baaij, 2018). CorA is a pentamer with a total of 10 TM helices, MgtE is a dimer with 10, and TRPM7 has 24. Our structure shows that 6 helices suffice to form a large cavity with ion binding sites deep within the membrane. The structure clearly demonstrates CNNMs are ion transporters rather than regulators of other proteins.

The specificity of the archaeal MtCNNM is unknown. The majority of studies of CNNM proteins have suggested that they transport Mg^2+^ ions. In *S. aureus*, MpfA likely functions to export Mg^2+^ as deletion mutants are unable to grow in the presence of high concentrations of Mg^2+^ (Armitano et al., 2016; Trachsel et al., 2019). Disruption of the homologous gene in *Bacillus subtilis* leads to increased cellular Mg^2+^ content, again supporting a role in Mg^2+^ efflux (Akanuma et al., 2014). In *Salmonella enterica*, a CNNM ortholog is required for reducing cytoplasmic concentration of the diamines, cadaverine and putrescine (Iwadate et al., 2021). In yeast, deletion of the CNNM ortholog, Mam3, confers resistance to high levels of manganese, cobalt and zinc (Yang et al., 2005). This could be an indirect, protective effect of elevated cytosolic Mg^2+^ levels; the Co^2+^ and Mn^2+^ resistance of CNNM mutants has been reported in multiple bacterial species including *S. aureus* (Armitano et al., 2016; Gibson et al., 1991; Pi et al., 2020).

Studies of the mammalian CNNMs have demonstrated an essential role in Mg^2+^ homeostasis. CNNM2 is essential in mice while loss of CNNM4 leads to male infertility and susceptibility to cancer (Funato et al., 2014; Yamazaki et al., 2019; Yamazaki et al., 2016). CNNM proteins act as tumor suppressors by depressing Mg^2+^-dependent energy metabolism and AMPK/mTOR signaling (Funato et al., 2014). CNNM2 and CNNM4 are abundant in the basolateral membranes of kidney and colon epithelial cells, where renal/intestinal (re)absorption of Mg^2+^ occurs (Stuiver et al., 2011; Yamazaki et al., 2013). Cellular studies have suggested that mammalian CNNMs act as a Na^+^/Mg^2+^ antiporters (Funato et al., 2014; Hirata et al., 2014; Yamazaki et al., 2013). Our observation of bound Mg^2+^ and Na^+^ ions in the MtCNNM structure is consistent with Na^+^/Mg^2+^ exchange driven transport. How this occurs is unknown, but one possibility is a rocker-switch mechanism with alternating access for ion binding on the two sides of the membrane (Fig. 5e).

The crystal structure of R235L MtCNNM provides insight into the regulatory function of the cytosolic domains. In humans, mutations that prevent Mg^2+^-ATP binding cause Jalili syndrome, hypomagnesemia, and block Mg^2+^ efflux in cellular assays (Chen et al., 2020). An analogous mutation, R251L, in *S. aureus* MpfA blocks protection against high concentrations of Mg^2+^ (Fig. 3g). Structurally, the loss of Mg^2+^-ATP binding disrupts dimerization of both the CBS-pair and AHB domains. The acidic residues in the AHB are reminiscent of the acidic patches in the cytosolic domains of CorA and MgtE that are involved in Mg^2+^ sensing and regulation of channel activity (Payandeh et al., 2013). Disruption of the AHB dimer could play a regulatory role through the loss of Mg^2+^ binding sites. Although not directly involved in Mg^2+^-ATP binding, the domains that follow the CBS-pair domains are also required for CNNM function. In human CNNM4, deletion of the CNBH domain decreases nucleotide binding and Mg^2+^ transport in cells likely due to a loss of CBS-pair dimerization (Chen et al., 2018). Deletion of the *S. aureus* MpfA CorC domain similarly blocks protection against Mg^2+^ (Fig. S5).

In animals, PRL phosphatases provide an additional level of regulation. PRL binding to CNNMs is regulated by PRL phosphorylation in response to Mg^2+^ levels (Gulerez et al., 2016; Kozlov et al., 2020). PRLs are highly oncogenic and promote cancer metastasis through their suppression of CNNM proteins (Funato et al., 2014; Yamazaki et al., 2019). The structures of a CNNM protein in both an active and inactive conformation provides a framework for understanding human CNNMs and the development of therapeutic approaches for regulating their activity.

## Methods

### Construction of phylogenetic tree

Amino acid sequences of various CNNM orthologs were aligned using MUSCLE (Edgar, 2004). The phylogenetic tree was generated using neighbor-joining method and bootstrapping of 1,000 replications in MEGAX (Version 10.1.8) (Kumar et al., 2018). The CNNM orthologs and their UniProt accession numbers are: cnnm2a (*Danio rerio*; A2ATX7), CNNM2 (*Homo sapiens*; Q9H8M5), CNNM4 (*Homo sapiens*; Q6P4Q7), CNNM4 (*Xenopus tropicalis*; A0JPA0), CNNM1 (*Homo sapiens*; Q9NRU3), CNNM3 (*Homo sapiens*; Q8NE01), UEX (*Drosophila melanogaster*; A0A0B7P9G0), cnnm-1 (*Caenorhabditis elegans*; A3QM97), MAM3 (*Saccharomyces cerevisiae*; Q12296), CBSDUF1 (*Arabidopsis thaliana*; Q67XQ0), CBSDUFCH1 (*Arabidopsis thaliana*; Q9LK65), MtCNNM (*Methanoculleus thermophilus*; A0A1G8XA46), CorB (*Salmonella typhimurium*; Q8XFY3), yfjD (*Escherichia coli*; P37908), MpfA (*Staphylococcus aureus*; A0A0H3JL60), and yhdP (*Bacillus subtilis*; O07585).

### Cloning of prokaryotic CNNMs

Codon-optimized cDNA of 20 prokaryotic CNNM orthologs were synthesized (Bio Basic Inc., Markham, Canada) and sub-cloned into NcoI and XhoI sites of pCGFP-BC vector(Kawate and Gouaux, 2006) with a C-terminal GFP-His8-tag for small-scale expression and detergent screening. Promising orthologs were subcloned into NdeI and XhoI sites of pET29a vector (Millipore Sigma) with a C-terminal His6-tag for large-scale expression and crystallization experiments. Constructs for *Methanoculleus thermophilus* CNNM (UniProt entry A0A1G8XA46): MtCNNM (residues 1-426), MtCNNMΔC (residues 1-322), MtCNNMΔC_Δloop_ (residues 1-322 Δ259-262), and MtCNNMΔC R235L. For constructs lacking TMD, MtCNNM_CBS+corc_ (residues 199-417), MtCNNM_CBS_ (residues 199-322), and MtCNNM_CorC_ (residues 333-417) were subcloned into BamHI and XhoI sites of pGEX-6P-1 vector (GE Healthcare) with an N-terminal GST-tag.

### Small-scale expression and detergent screening of prokaryotic CNNMs

Prokaryotic CNNMs cloned in pCGFP-BC were transformed into *E. coli* strain C41 (DE3). Cells were grown in Luria Broth (LB) at 37°C to an optical density of 0.6 and induced with 1 mM IPTG for 4 hours at 30°C. Cell pellet was obtained by centrifuging at 5,000 g for 10 min. The pellet was re-suspended in lysis buffer (50 mM HEPES, 500 mM NaCl, 5% glycerol, pH 7.5) supplemented with cOmplete^™^ protease inhibitor cocktail (Roche) and split into 6 fractions. Lysis was performed using a 24-probe sonicator. Each fraction was solubilized with a different detergent (DDM; LMNG; OGNG; LDAO; C12E9; OG) to final concentration of 1%, then purified by IMAC in the same detergent (3x CMC). Elutions were analyzed by SDS-PAGE and size-exclusion chromatography on SEPAX Zenic-C SEC-300 connected to fluorescence detector (Ex 480 nm/ Em 510 nm) in buffer (50 mM HEPES pH 7.5, 150 mM NaCl, 0.5 mM TCEP, 3x CMC of detergent).

### Expression and purification of MtCNNMandMtCNNMΔC

Constructs were transformed into *E. coli* strain C41 (DE3). Cells were grown in Luria Broth (LB) at 37°C to an optical density of 0.6 and induced with 0.5 mM IPTG overnight at 18°C. Cell pellet was obtained by centrifuging at 5,000 g for 20 min. The pellet was re-suspended in lysis buffer (50 mM HEPES, 500 mM NaCl, 5% glycerol, pH 7.5) supplemented with cOmplete^™^ protease inhibitor cocktail (Roche), 10 μg/mL DNAse I, 1 mM CaCl2, 0.1 mg/mL lysozyme. Cells were lysed by passing through Avestin Emulsiflex-C3 homogenizer (10,000 – 15,000 p. s.i.). Cellular debris were removed by centrifugation at 27,000 g for 10 min at 4°C (this step is omitted for fulllength MtCNNM). Membranes were pelleted by ultracentrifugation at 150,000 g for 1 hour at 4°C. The membrane fraction was collected, flash frozen in liquid nitrogen, and stored at −80°C for later use. The membrane fraction after thawing was solubilized in lysis buffer supplemented with 1% DDM for 1 hour at 4°C on a rotator, then ultracentrifuged at 150,000 g for 30 min at 4°C. The supernatant was loaded onto Qiagen Ni-NTA resin by batch binding, and incubated with gentle shaking for 1 hour at 4°C. The resin was then washed with wash buffer (50 mM HEPES, 500 mM NaCl, 5% glycerol, 30 mM imidazole, pH 7.5) containing 0.03% DDM or 0.05% UDM and eluted with elution buffer (50 mM HEPES, 500 mM NaCl, 5% glycerol, 300 mM imidazole, pH 7.5) containing 0.03% DDM or 0.05% UDM. The affinity-purified protein was further purified by size exclusion chromatography on a HiLoad 16/600 Superdex 200 pg column (GE Healthcare) in HPLC buffer (20 mM HEPES, 150 mM NaCl, pH 7.5) containing 0.03% DDM or 0.05% UDM. The final purified proteins were concentrated using 50 kDa (MtCNNMΔC) or 100 kDa (MtCNNM) cutoff concentrators (Amicon Ultra, Millipore), flash frozen in liquid nitrogen, and stored at −80°C for later use. The protein concentration is determined spectrophotometrically using Nanodrop, and purity is verified by SDS-PAGE.

### Expression and purification of MtCNNM constructs lacking TMD

Constructs were transformed into *E. coli* strain BL21 (DE3). Cells were grown in Luria Broth (LB) at 37°C to an optical density of 0.8 and induced with 1 mM IPTG overnight at 20°C. Cell pellet was obtained by centrifuging at 5,000 g for 20 min. The pellet was re-suspended in lysis buffer (50 mM HEPES, 500 mM NaCl, 5% glycerol, pH 7.5) supplemented with cOmplete^™^ protease inhibitor cocktail (Roche), 10 μg/mL DNAse I, 1 mM CaCl2, and 0.1 mg/mL lysozyme. Cells were lysed by sonication. Cellular debris were removed by centrifugation at 44,000 g for 45 min at 4°C. The supernatant was loaded onto Glutathione Sepharose resin (GE Healthcare), washed with lysis buffer and eluted with lysis buffer containing 20 mM glutathione. The GST-tag was removed by overnight incubation with PreScission Protease, leaving an N-terminal Gly-Pro-Leu-Gly-Ser extension. The protein was further purified by size exclusion chromatography on a HiLoad 16/600 Superdex 75 pg column (GE Healthcare) in HPLC buffer (20 mM HEPES, 100 mM NaCl, pH 7.5). The protein was diluted to 5 μM, dialyzed overnight in dialysis buffer (20 mM HEPES, 100 mM NaCl, 5 mM EDTA, pH 7.5), and re-injected onto Superdex-75 to remove bound nucleotides. The final purified protein was concentrated to around 10 mg/mL (measured by NanoDrop), and the purity verified by SDS-PAGE.

### Crystallization

Crystals of MtCNNM_CBS_ co-crystallized with 5 mM ATP were obtained by equilibrating 0.4 μL of protein (20.8 mg/mL MtCNNMΔC purified in 0.03% DDM) and 0.4 μL of reservoir solution (0.1 M MOPS, pH 7.0; 9% PEG 8000; 20 mM MgCl_2_) in sitting-drop vapor diffusion system incubated at 22°C. Rod-like crystals appeared after 2 weeks. The crystal was cryo-protected with reservoir solution supplemented with 5 mM ATP and 30% ethylene glycol, picked up in a nylon loop, and flash-cooled in a N_2_ cold stream.

Crystals of MtCNNMΔC_Δloop_ co-crystallized with 5 mM ATP were obtained by equilibrating 1 μL of protein (19.5 mg/mL MtCNNMΔC_Δloop_ purified in 0.05% UDM) and 1 μL of reservoir solution (0.1 M sodium citrate, pH 5.5; 0.1 M Li2SO4; 0.1 M NaCl; 20 mM MgCl_2_; 34% PEG400; 10 mM Na_2_HPO_4_) in hanging-drop vapor diffusion system incubated at 22°C. Petal-like crystals appear after 1 week. The crystal was directly picked up in a nylon loop and flash-cooled in a N_2_ cold stream.

Crystals of MtCNNMΔC R235L were obtained by equilibrating 1 μL of protein (4.6 mg/mL MtCNNMΔC R235L purified in HPLC buffer containing 20 mM HEPES, 500 mM NaCl, 0.05% UDM, pH 7.5) and 1 μL of reservoir solution (0.1 M sodium citrate, pH 5.5; 0.1 M Li2SO4; 13.5% PEG4000) in hanging-drop vapor diffusion system incubated at 22°C. Rod-like crystals appear after 1 day. The crystal was cryo-protected with reservoir solution supplemented with 20% glycerol, picked up in a nylon loop, and flash-cooled in a N_2_ cold stream.

### Data collection and structure determination

The MtCNNM_CBS_ dataset from a single crystal was collected using a Pilatus3 6M detector at beamline 5.0.2 of Advanced Light Source (ALS). Data processing and scaling were performed with HKL-2000 (Otwinowski and Minor, 1997) with auto-corrections enabled. Initial phases were obtained by molecular replacement with Phaser (McCoy et al., 2007) in PHENIX (Adams et al., 2010) using CBS-pair domain structure of CorC (PDB: 5YZ2) (Feng et al., 2018). The model was subsequently improved through iterative cycles of manual building with Coot (Emsley et al., 2010) and refinement with phenix.refine (Afonine et al., 2012). TLS parameters were included at later stages of the refinement (Painter and Merritt, 2006).

The MtCNNMΔC_Δloop_ dataset from a single crystal was collected using a Pilatus3 6M detector at beamline 08ID-1 of the Canadian Macromolecular Crystallography Facility (CMCF) of the Canadian Light Source (CLS). The dataset showed anisotropic diffraction up to 3.25 Å. Data processing and scaling were performed with HKL-2000 (Otwinowski and Minor, 1997) with auto-corrections enabled, in which ellipsoid truncation was performed automatically. Resolution cut-off is based on CC1/2 = 0.3 (Karplus and Diederichs, 2015). Resolution limits after ellipsoid truncation were a*= 4.07 Å, b* = 3.71 Å and c* = 3.25 Å. Initial phases for the CBS-pair domain were obtained by molecular replacement with Phaser (McCoy et al., 2007) in PHENIX (Adams et al., 2010) using MtCNNM_CBS_ structure (determined in this study). Then AutoBuild (Terwilliger et al., 2008) was used to build in the missing domains (TMD and AHB). The model was then improved through iterative cycles of manual building with Coot (Emsley et al., 2010) and refinement with phenix.refine (Afonine et al., 2012). TLS parameters were included at later stages of the refinement (Painter and Merritt, 2006).

The MtCNNMΔC R235L dataset from a single crystal was collected using Pilatus3 6M detector at beamline 5.0.2 of Advanced Light Source (ALS). The dataset was processed with DIALS (Winter et al., 2018) and scaled with Aimless (Evans and Murshudov, 2013). Initial phases were obtained by molecular replacement with Phaser (McCoy et al., 2007) in PHENIX (Adams et al., 2010) using the CBS-pair and TMD structures (determined in this study). The model was subsequently improved through iterative cycles of manual building with Coot (Emsley et al., 2010) and refinement with phenix.refine (Afonine et al., 2012).

The final structures were validated with MolProbity (Chen et al., 2010). Crystallographic data collection and structure refinement statistics are shown in Table S1. Structural images were prepared with PyMOL, Version 2.3.4 (Schrödinger LLC, New York). Electrostatic surface potentials were calculated using the APBS plugin within PyMOL (Jurrus et al., 2018).

### In vivo complementation assays of various MpfA mutants in Staphylococcus aureus

Various point mutants of mpfA were cloned on a multicopy plasmid (pCN47 based) under the control of a xylose inducible promoter (Charpentier et al., 2004; Tu Quoc et al., 2007). All mutated alleles were obtained by fusion PCR and cloned between restrictions sites SphI and AscI.

The functionality of the alleles was assessed by testing their ability to complement the magnesium sensitivity of an *Staphylococcus aureus* ΔmpfA strain (PR01-36) (Armitano et al., 2016). Overnight cultures of strains carrying various plasmids were serially diluted in Mueller Hinton (MH) media and 10 μL of each dilution were spotted onto MH plates containing media (MH 211443, BD Biosciences, Allschwil, Switzerland) supplemented with 10 mg/L uracil, 10 mg/L erythromycin, 13 g/L of agar and as necessary, varying concentrations of MgCl_2_ and xylose. Plates were incubated for 20 hours at 37°C. Only relevant dilutions (10^-4^ to 10^-6^) are shown in figures. All experiments include three controls: WT strain (PR01) carrying an empty vector, ΔmpfA strain carrying an empty vector, and ΔmpfA strain carrying a vector containing mpfA G326C (an inactive allele).

### Isothermal titration calorimetry

ITC experiments were performed on a MicroCal VP-ITC titration calorimeter (Malvern Instruments Ltd) at 20°C with stirring at 310 rpm. TMD-containing constructs (15 μM final concentration) and ligands were prepared in HPLC buffer containing 0.05% UDM (± 50 mM MgCl_2_). Cytosolic constructs (30 μM final concentration) were prepared in HPLC buffer (± 10 or 50 mM MgCl_2_). The ligands were injected 19 times (5 μL for the first injection, 15 μL for subsequent injections), with 4 min intervals between injections. Results were analyzed using ORIGIN software (MicroCal) and fitted to a binding model with a single set of identical sites.

### Analytical ultracentrifugation

Sedimentation velocity AUC experiments were performed at 20°C using a Beckman Coulter XL-I Optima analytical ultracentrifuge and an An-60Ti rotor at 98,000 g (35,000 rpm) for 18 hours with scans performed every 60 seconds. A double-sector cell, equipped with a 12 mm Epon centerpiece and sapphire windows, was loaded with 380 and 400 μL of sample and HPLC buffer. MtCNNM_CBS_ (100 μM) with various ligands were monitored using interference optics. The data were analyzed with Sedfit v1501b (Schuck, 2000) using a continuous c(s) distribution. Numerical values for the solvent density, viscosity, and partial specific volume were determined using Sednterp (Laue et al., 1992). Buffer density and viscosity were calculated to be 1.0039 g/cm^3^ and 0.01026 mPa·s, respectively (20 mM HEPES, 100 mM NaCl, pH 7.5). Partial specific volumes for MtCNNM_CBS_ was calculated to be 0.7464 cm^3^/g. The frictional ratio (f/f_0_) value for MtCNNM_CBS_ was calculated using US-SOMO (Brookes et al., 2010) to be 1.26. Residual and c(s) distribution graphs were plotted using GUSSI (Brautigam, 2015).

### Production and purification of MSP1D1

pMSP1D1 was a gift from S. Sligar (Addgene plasmid 20061). MSP1D1 production was carried out according to published protocols (Ritchie et al., 2009). In brief, *E. coli* BL21 (DE3) cells transformed with pMSP1D1 were grown in LB at 37°C to an optical density of 0.8 and induced with 1 mM IPTG for 4 hours at 30°C. MSP1D1 was purified by nickel affinity chromatography according to standard conditions described in (Ritchie et al., 2009). The polyhistidine tag was removed by overnight incubation with TEV protease and further purified by Superdex-75 size-exclusion column (GE Healthcare) in HPLC buffer (20 mM HEPES, 150 mM NaCl, pH 7.5)

### Reconstitution into nanodisc

MtCNNM and MSP1D1 were mixed with soybean polar extract (Avanti) solubilized in 40 mM DDM at a MtCNNM:MSP1D1:lipid molar ratio of 2:10:550 in HPLC buffer. Detergent was removed by incubation with Bio-Beads (Bio-Rad SM-2 Resin) at 4°C overnight with constant rotation. Bio-beads were removed via filtration and the reconstitution mixture was re-loaded onto Qiagen Ni-NTA resin to remove empty nanodiscs. The resin was washed with wash buffer (20 mM HEPES, 200 mM NaCl, 20 mM imidazole, pH 7.5) and eluted with wash buffer with 300 mM imidazole. The eluted protein was further purified by size exclusion chromatography on a Superdex 200 Increase 10/300 GL column (GE Healthcare) in HPLC buffer (20 mM HEPES, 150 mM NaCl, pH 7.5).

### Hydrogen deuterium exchange mass spectrometry

HDX-MS reactions were performed in a similar manner as described previously (Jenkins et al., 2018; Lucic et al., 2018). In brief, HDX reactions for MtCNNM and MtCNNMΔC were conducted in a final reaction volume of 10 μL with a molar quantity of 20 pmol of MtCNNM and MtCNNMΔC. The reaction was started by the addition of 9.0 μL of D_2_O buffer (100 mM NaCl, 20 mM HEPES pH 7.5, 94% D_2_O (V/V)) to 1.0 μL of protein solution (final D_2_O concentration of 84.9%). The reaction proceeded for 3, 30, 300, or 3000 s at 20°C, before being quenched with ice cold acidic quench buffer, resulting in a final concentration of 0.6 M guanidine-HCl and 0.9% formic acid post quench. All conditions and timepoints were created and run in triplicate. Samples were flash frozen immediately after quenching and stored at −80°C until injected onto the ultra-performance liquid chromatography (UPLC) system for proteolytic cleavage, peptide separation, and injection onto a QTOF for mass analysis, described below.

Protein samples were rapidly thawed and injected onto a UPLC system kept in a Peltier driven cold box at 2°C (LEAP). The protein was run over two immobilized pepsin columns (Trajan; ProDx Protease column PDX.PP01-F32) and the peptides were collected onto a VanGuard Precolumn trap (Waters). The trap was eluted in line with an ACQUITY 1.7 μm particle, 100 × 1 mm^2^ C18 UPLC column (Waters), using a gradient of 5%-36% B (Buffer A 0.1% formic acid, Buffer B 100% acetonitrile) over 16 min. MS experiments were performed on an Impact HD QTOF (Bruker) and peptide identification was done by running tandem MS (MS/MS) experiments run in data-dependent acquisition mode. The resulting MS/MS datasets were analyzed using PEAKS7 (PEAKS) and a false discovery rate was set at 1% using a database of purified proteins and known contaminants. HDExaminer Software (Sierra Analytics) was used to automatically calculate the level of deuterium incorporation into each peptide. All peptides were manually inspected for correct charge state and presence of overlapping peptides.

Deuteration levels were calculated using the centroid of the experimental isotope clusters. Differences in exchange in a peptide were considered significant if they met three of the following criteria: > 5% change in exchange, > 0.4 Da mass difference in exchange, a p-value < 0.01 using a two-tailed Student’s t-test, and change spanned by multiple peptides.

#### Homology modeling of human CNNM2

The homology model of human CNNM2 was constructed with MODELLER Version 9.25 (Webb and Sali, 2016) using MtCNNMΔC structure as the template. The sequence alignment was performed using Clustal Omega (Madeira et al., 2019). The model was evaluated using PROCHECK (Laskowski et al., 1993). Conservation coloring was done with ProtSkin (http://www.mcgnmr.mcgill.ca/ProtSkin/).

#### Molecular dynamics simulation

MtCNNMΔC structure was used as the starting model for MD simulations. The transporter was embedded in a bilayer of 3POPC: 1POPG lipids and solvated in 150 mM NaCl (neutral with 277 Na^+^ and 139^-^ Cl ions) using the web service CHARMM-GUI (Jo et al., 2008; Wu et al., 2014). Most residues were assigned their standard protonation state at pH 7. The total number of atoms in the atomic model is on the order of 200,000. The all-atom CHARMM force field PARAM36 for protein (Best et al., 2012; MacKerell et al., 1998; MacKerell et al., 2004), lipids (Klauda et al., 2010), and ions (Beglov and Roux, 1994) was used. Explicit water was represented with the TIP3P model (Jorgensen et al., 1983). The model was refined using energy minimization for at least 2,000 steps. All the simulations were performed under NPT (constant number of particle N, pressure P, and temperature T) conditions at 303 K and 1 atm, and periodic boundary conditions with electrostatic interactions were treated by the particle mesh Ewald method (Darden et al., 1993) and a real-space cutoff of 12 Å. The simulations use a time step of 2 fs, with bond distances involving hydrogen atoms fixed using the SHAKE algorithm (Ryckaert et al., 1977). After minimization, positional restraints on all of the Cα atoms were gradually released after which a trajectory of 500 ns was generated using NAMD version 2.11 (Phillips et al., 2005).

#### Targeted molecular dynamics simulation

For preparing the target (open) model, the interface helices of MtCNNMΔC were pulled upward around the region of Glu111, causing the TMD to open. All forces were then removed and the whole system was equilibrated for ~ 8 ns. For the targeted MD simulation, the MtCNNMΔC structure was used as the starting model. During the simulation, forces were applied on the backbone of the transmembrane helices with a force constant 200 kcal·mol^-1^·A^-2^ for a period of 10 ns. This decreases the RMSD value linearly to reach the final target conformation.

## Data availability

Atomic coordinates and structure factors for MtCNNM_CBS_ with Mg^2+^-ATP, MtCNNMΔC with Mg^2+^-ATP, and MtCNNMΔC R235L mutant have been deposited to the Protein Data Bank (www.rcsb.org) under accession numbers 7LJ6, 7LJ7, and 7LJ8, respectively.

## Acknowledgements

We thank Katharina Duerr and her lab members for assistance with initial screening of prokaryotic CNNMs; Alexei Gorelik for advice on crystallographic data processing; Martin Schmeing and Camille Fortinez for crystallographic data collection at Advanced Light Source (ALS); Canadian Light Source (CLS) staff: Michel Fodje, Shaun Labiuk, Bulat Gabidullin, and Kathyrn Janzen. CLS is supported by the Canada Foundation for Innovation, the Natural Sciences and Engineering Research Council (NSERC), the National Research Council, the Canadian Institutes of Health Research (CIHR), the Government of Saskatchewan, and the University of Saskatchewan. Y.S.C. is supported by a CIHR Doctoral Research Award (GSD-167011). This work is supported by NSERC grant RGPIN-2020-07195 to K.G.

## Author Contributions

Y.S.C. designed experiments, cloned constructs, performed small-scale screenings, purified proteins, and solved crystal structures. G.K. assisted with crystal screening and performed ITC experiments. J.A. performed complementation assays. R.F. performed AUC and ITC experiments. B.E.M. performed HDX-MS experiments. A.R. performed molecular dynamics simulations. Y.S.C. and K.G. wrote the manuscript. K.G., J.E.B, and B.R. oversaw the research.

## References

Adams, P.D., Afonine, P.V., Bunkoczi, G., Chen, V.B., Davis, I.W., Echols, N., Headd, J.J., Hung, L.W., Kapral, G.J., Grosse-Kunstleve, R.W., et al. (2010). PHENIX: a comprehensive Python-based system for macromolecular structure solution. Acta Crystallogr D Biol Crystallogr 66, 213–221.

Afonine, P.V., Grosse-Kunstleve, R.W., Echols, N., Headd, J.J., Moriarty, N.W., Mustyakimov, M., Terwilliger, T.C., Urzhumtsev, A., Zwart, P.H., and Adams, P.D. (2012). Towards automated crystallographic structure refinement with phenix.refine. Acta Crystallogr D Biol Crystallogr 68, 352–367.

Akanuma, G., Kobayashi, A., Suzuki, S., Kawamura, F., Shiwa, Y., Watanabe, S., Yoshikawa, H., Hanai, R., and Ishizuka, M. (2014). Defect in the formation of 70S ribosomes caused by lack of ribosomal protein L34 can be suppressed by magnesium. J Bacteriol 196, 3820–3830.

Arjona, F.J., de Baaij, J.H., Schlingmann, K.P., Lameris, A.L., van Wijk, E., Flik, G., Regele, S., Korenke, G.C., Neophytou, B., Rust, S., et al. (2014). CNNM2 mutations cause impaired brain development and seizures in patients with hypomagnesemia. PLoS Genet 10, e1004267.

Arjona, F.J., and de Baaij, J.H.F. (2018). CrossTalk opposing view: CNNM proteins are not Na(+) /Mg(2+) exchangers but Mg(2+) transport regulators playing a central role in transepithelial Mg(2+) (re)absorption. J Physiol 596, 747–750.

Armitano, J., Redder, P., Guimaraes, V.A., and Linder, P. (2016). An Essential Factor for High Mg(2+) Tolerance of Staphylococcus aureus. Front Microbiol 7, 1888.

Beglov, D., and Roux, B. (1994). Finite representation of an infinite bulk system: solvent boundary potential for computer simulations. The Journal of chemical physics 100, 9050–9063.

Best, R.B., Zhu, X., Shim, J., Lopes, P.E., Mittal, J., Feig, M., and Mackerell, A.D., Jr. (2012). Optimization of the additive CHARMM all-atom protein force field targeting improved sampling of the backbone phi, psi and side-chain chi(1) and chi(2) dihedral angles. J Chem Theory Comput 8, 3257–3273.

Blum, M., Chang, H.Y., Chuguransky, S., Grego, T., Kandasaamy, S., Mitchell, A., Nuka, G., Paysan-Lafosse, T., Qureshi, M., Raj, S., et al. (2021). The InterPro protein families and domains database: 20 years on. Nucleic Acids Res 49, D344–D354.

Brautigam, C.A. (2015). Calculations and Publication-Quality Illustrations for Analytical Ultracentrifugation Data. Methods Enzymol 562, 109–133.

Brookes, E., Demeler, B., and Rocco, M. (2010). Developments in the US-SOMO bead modeling suite: new features in the direct residue-to-bead method, improved grid routines, and influence of accessible surface area screening. Macromol Biosci 10, 746–753.

Charpentier, E., Anton, A.I., Barry, P., Alfonso, B., Fang, Y., and Novick, R.P. (2004). Novel cassette-based shuttle vector system for gram-positive bacteria. Appl Environ Microbiol 70, 6076–6085.

Chen, V.B., Arendall, W.B., 3rd, Headd, J.J., Keedy, D.A., Immormino, R.M., Kapral, G.J., Murray, L.W., Richardson, J.S., and Richardson, D.C. (2010). MolProbity: all-atom structure validation for macromolecular crystallography. Acta Crystallogr D Biol Crystallogr 66, 12–21.

Chen, Y.S., Kozlov, G., Fakih, R., Funato, Y., Miki, H., and Gehring, K. (2018). The cyclic nucleotide-binding homology domain of the integral membrane protein CNNM mediates dimerization and is required for Mg(2+) efflux activity. J Biol Chem 293, 19998–20007.

Chen, Y.S., Kozlov, G., Fakih, R., Yang, M., Zhang, Z., Kovrigin, E.L., and Gehring, K. (2020). Mg(2+)-ATP Sensing in CNNM, a Putative Magnesium Transporter. Structure 28, 324–335 e324.

Corral-Rodriguez, M.A., Stuiver, M., Abascal-Palacios, G., Diercks, T., Oyenarte, I., Ereno-Orbea, J., de Opakua, A.I., Blanco, F.J., Encinar, J.A., Spiwok, V., et al. (2014). Nucleotide binding triggers a conformational change of the CBS module of the magnesium transporter CNNM2 from a twisted towards a flat structure. Biochem J 464, 23–34.

Daneshmandpour, Y., Darvish, H., Pashazadeh, F., and Emamalizadeh, B. (2019). Features, genetics and their correlation in Jalili syndrome: a systematic review. J Med Genet 56, 358–369.

Darden, T., York, D., and Pedersen, L. (1993). Particle mesh Ewald: An No log (N) method for Ewald sums in large systems. The Journal of chemical physics 98, 10089–10092.

de Baaij, J.H., Hoenderop, J.G., and Bindels, R.J. (2015). Magnesium in man: implications for health and disease. Physiol Rev 95, 1–46.

de Baaij, J.H., Stuiver, M., Meij, I.C., Lainez, S., Kopplin, K., Venselaar, H., Muller, D., Bindels, R.J., and Hoenderop, J.G. (2012). Membrane topology and intracellular processing of cyclin M2 (CNNM2). J Biol Chem 287, 13644–13655.

Duan, J., Li, Z., Li, J., Hulse, R.E., Santa-Cruz, A., Valinsky, W.C., Abiria, S.A., Krapivinsky, G., Zhang, J., and Clapham, D.E. (2018). Structure of the mammalian TRPM7, a magnesium channel required during embryonic development. Proc Natl Acad Sci U S A 115, E8201–E8210.

Edgar, R.C. (2004). MUSCLE: multiple sequence alignment with high accuracy and high throughput. Nucleic Acids Res 32, 1792–1797.

El-Gebali, S., Mistry, J., Bateman, A., Eddy, S.R., Luciani, A., Potter, S.C., Qureshi, M., Richardson, L.J., Salazar, G.A., Smart, A., et al. (2019). The Pfam protein families database in 2019. Nucleic Acids Res 47, D427–D432.

Emsley, P., Lohkamp, B., Scott, W.G., and Cowtan, K. (2010). Features and development of Coot. Acta Crystallogr D Biol Crystallogr 66, 486–501.

Ereno-Orbea, J., Oyenarte, I., and Martinez-Cruz, L.A. (2013). CBS domains: Ligand binding sites and conformational variability. Arch Biochem Biophys 540, 70–81.

Eshaghi, S., Niegowski, D., Kohl, A., Martinez Molina, D., Lesley, S.A., and Nordlund, P. (2006). Crystal structure of a divalent metal ion transporter CorA at 2.9 angstrom resolution. Science 313, 354–357.

Evans, P.R., and Murshudov, G.N. (2013). How good are my data and what is the resolution? Acta Crystallogr D Biol Crystallogr 69, 1204–1214.

Feng, N., Qi, C., Hou, Y.J., Zhang, Y., Wang, D.C., and Li, D.F. (2018). The C2’- and C3’-endo equilibrium for AMP molecules bound in the cystathionine-beta-synthase domain. Biochem Biophys Res Commun 497, 646–651.

Funato, Y., Furutani, K., Kurachi, Y., and Miki, H. (2018). CrossTalk proposal: CNNM proteins are Na(+) /Mg(2+) exchangers playing a central role in transepithelial Mg(2+) (re)absorption. J Physiol 596, 743–746.

Funato, Y., and Miki, H. (2019). Molecular function and biological importance of CNNM family Mg2+ transporters. J Biochem 165, 219–225.

Funato, Y., Yamazaki, D., Mizukami, S., Du, L., Kikuchi, K., and Miki, H. (2014). Membrane protein CNNM4-dependent Mg2+ efflux suppresses tumor progression. J Clin Invest 124, 5398–5410.

Gibson, M.M., Bagga, D.A., Miller, C.G., and Maguire, M.E. (1991). Magnesium transport in Salmonella typhimurium: the influence of new mutations conferring Co2+ resistance on the CorA Mg2+ transport system. Mol Microbiol 5, 2753–2762.

Gulerez, I., Funato, Y., Wu, H., Yang, M., Kozlov, G., Miki, H., and Gehring, K. (2016). Phosphocysteine in the PRL-CNNM pathway mediates magnesium homeostasis. EMBO Rep 17, 1890–1900.

Hardy, S., Uetani, N., Wong, N., Kostantin, E., Labbe, D.P., Begin, L.R., Mes-Masson, A., Miranda-Saavedra, D., and Tremblay, M.L. (2015). The protein tyrosine phosphatase PRL-2 interacts with the magnesium transporter CNNM3 to promote oncogenesis. Oncogene 34, 986–995.

Hattori, M., Tanaka, Y., Fukai, S., Ishitani, R., and Nureki, O. (2007). Crystal structure of the MgtE Mg2+ transporter. Nature 448, 1072–1075.

Hirata, Y., Funato, Y., Takano, Y., and Miki, H. (2014). Mg2+-dependent interactions of ATP with the cystathionine-beta-synthase (CBS) domains of a magnesium transporter. J Biol Chem 289, 14731–14739.

Iwadate, Y., Ramezanifard, R., Golubeva, Y.A., Fenlon, L.A., and Slauch, J.M. (2021). PaeA (YtfL) protects from cadaverine and putrescine stress in Salmonella Typhimurium and E. coli. Mol Microbiol.

Jenkins, M.L., Margaria, J.P., Stariha, J.T.B., Hoffmann, R.M., McPhail, J.A., Hamelin, D.J., Boulanger, M.J., Hirsch, E., and Burke, J.E. (2018). Structural determinants of Rab11 activation by the guanine nucleotide exchange factor SH3BP5. Nat Commun 9, 3772.

Jo, S., Kim, T., Iyer, V.G., and Im, W. (2008). CHARMM-GUI: a web-based graphical user interface for CHARMM. J Comput Chem 29, 1859–1865.

Jorgensen, W.L., Chandrasekhar, J., Madura, J.D., Impey, R.W., and Klein, M.L. (1983). Comparison of simple potential functions for simulating liquid water. The Journal of chemical physics 79, 926–935.

Jurrus, E., Engel, D., Star, K., Monson, K., Brandi, J., Felberg, L.E., Brookes, D.H., Wilson, L., Chen, J., Liles, K., et al. (2018). Improvements to the APBS biomolecular solvation software suite. Protein Sci 27, 112–128.

Karplus, P.A., and Diederichs, K. (2015). Assessing and maximizing data quality in macromolecular crystallography. Curr Opin Struct Biol 34, 60–68.

Kawate, T., and Gouaux, E. (2006). Fluorescence-detection size-exclusion chromatography for precrystallization screening of integral membrane proteins. Structure 14, 673–681.

Klauda, J.B., Venable, R.M., Freites, J.A., O’Connor, J.W., Tobias, D.J., Mondragon-Ramirez, C., Vorobyov, I., MacKerell, A.D., Jr., and Pastor, R.W. (2010). Update of the CHARMM all-atom additive force field for lipids: validation on six lipid types. J Phys Chem B 114, 7830–7843.

Kozlov, G., Funato, Y., Chen, Y.S., Zhang, Z., Illes, K., Miki, H., and Gehring, K. (2020). PRL3 pseudophosphatase activity is necessary and sufficient to promote metastatic growth. J Biol Chem 295, 11682–11692.

Kumar, S., Stecher, G., Li, M., Knyaz, C., and Tamura, K. (2018). MEGA X: Molecular Evolutionary Genetics Analysis across Computing Platforms. Mol Biol Evol 35, 1547–1549.

Laskowski, R.A., MacArthur, M.W., Moss, D.S., and Thornton, J.M. (1993). PROCHECK: a program to check the stereochemical quality of protein structures. Journal of applied crystallography 26, 283–291.

Laue, T., Bhairavi, D., Ridgeway, T., and Pelletier, S. (1992). SE Harding, and AJ Rowe, and JC Horton, eds. Analytical Ultracentrifugation in Biochemistry and Polymers Science 90–125 (Royal Society of Chemistry, Cambridge, UK).

Lucic, I., Rathinaswamy, M.K., Truebestein, L., Hamelin, D.J., Burke, J.E., and Leonard, T.A. (2018). Conformational sampling of membranes by Akt controls its activation and inactivation. Proc Natl Acad Sci U S A 115, E3940–E3949.

Lunin, V.V., Dobrovetsky, E., Khutoreskaya, G., Zhang, R., Joachimiak, A., Doyle, D.A., Bochkarev, A., Maguire, M.E., Edwards, A.M., and Koth, C.M. (2006). Crystal structure of the CorA Mg2+ transporter. Nature 440, 833–837.

MacKerell, A.D., Bashford, D., Bellott, M., Dunbrack, R.L., Evanseck, J.D., Field, M.J., Fischer, S., Gao, J., Guo, H., Ha, S., et al. (1998). All-atom empirical potential for molecular modeling and dynamics studies of proteins. J Phys Chem B 102, 3586–3616.

MacKerell, A.D., Jr., Feig, M., and Brooks, C.L., 3rd (2004). Improved treatment of the protein backbone in empirical force fields. J Am Chem Soc 126, 698–699.

Madeira, F., Park, Y.M., Lee, J., Buso, N., Gur, T., Madhusoodanan, N., Basutkar, P., Tivey, A.R.N., Potter, S.C., Finn, R.D., et al. (2019). The EMBL-EBI search and sequence analysis tools APIs in 2019. Nucleic Acids Res 47, W636–W641.

McCoy, A.J., Grosse-Kunstleve, R.W., Adams, P.D., Winn, M.D., Storoni, L.C., and Read, R.J. (2007). Phaser crystallographic software. J Appl Crystallogr 40, 658–674.

Otwinowski, Z., and Minor, W. (1997). [20] Processing of X-ray diffraction data collected in oscillation mode. Methods Enzymol 276, 307–326.

Painter, J., and Merritt, E.A. (2006). Optimal description of a protein structure in terms of multiple groups undergoing TLS motion. Acta Crystallogr D Biol Crystallogr 62, 439–450.

Parveen, A., Mirza, M.U., Vanmeert, M., Akhtar, J., Bashir, H., Khan, S., Shehzad, S., Froeyen, M., Ahmed, W., Ansar, M., et al. (2019). A novel pathogenic missense variant in CNNM4 underlying Jalili syndrome: Insights from molecular dynamics simulations. Mol Genet Genomic Med 7, e902.

Payandeh, J., and Pai, E.F. (2006). A structural basis for Mg2+ homeostasis and the CorA translocation cycle. EMBO J 25, 3762–3773.

Payandeh, J., Pfoh, R., and Pai, E.F. (2013). The structure and regulation of magnesium selective ion channels. Biochim Biophys Acta 1828, 2778–2792.

Phillips, J.C., Braun, R., Wang, W., Gumbart, J., Tajkhorshid, E., Villa, E., Chipot, C., Skeel, R.D., Kale, L., and Schulten, K. (2005). Scalable molecular dynamics with NAMD. J Comput Chem 26, 1781–1802.

Pi, H., Wendel, B.M., and Helmann, J.D. (2020). Dysregulation of magnesium transport protects Bacillus subtilis against manganese and cobalt intoxication. J Bacteriol.

Ritchie, T.K., Grinkova, Y.V., Bayburt, T.H., Denisov, I.G., Zolnerciks, J.K., Atkins, W.M., and Sligar, S.G. (2009). Chapter 11 - Reconstitution of membrane proteins in phospholipid bilayer nanodiscs. Methods Enzymol 464, 211–231.

Ritter, B., Denisov, A.Y., Philie, J., Deprez, C., Tung, E.C., Gehring, K., and McPherson, P.S. (2004). Two WXXF-based motifs in NECAPs define the specificity of accessory protein binding to AP-1 and AP-2. EMBO J 23, 3701–3710.

Ryckaert, J.-P., Ciccotti, G., and Berendsen, H.J. (1977). Numerical integration of the cartesian equations of motion of a system with constraints: molecular dynamics of n-alkanes. Journal of computational physics 23, 327–341.

Schuck, P. (2000). Size-distribution analysis of macromolecules by sedimentation velocity ultracentrifugation and lamm equation modeling. Biophys J 78, 1606–1619.

Stuiver, M., Lainez, S., Will, C., Terryn, S., Gunzel, D., Debaix, H., Sommer, K., Kopplin, K., Thumfart, J., Kampik, N.B., et al. (2011). CNNM2, encoding a basolateral protein required for renal Mg2+ handling, is mutated in dominant hypomagnesemia. Am J Hum Genet 88, 333–343.

Terwilliger, T.C., Grosse-Kunstleve, R.W., Afonine, P.V., Moriarty, N.W., Zwart, P.H., Hung, L.W., Read, R.J., and Adams, P.D. (2008). Iterative model building, structure refinement and density modification with the PHENIX AutoBuild wizard. Acta Crystallogr D Biol Crystallogr 64, 61–69.

Trachsel, E., Redder, P., Linder, P., and Armitano, J. (2019). Genetic screens reveal novel major and minor players in magnesium homeostasis of Staphylococcus aureus. PLoS Genet 15, e1008336.

Tu Quoc, P.H., Genevaux, P., Pajunen, M., Savilahti, H., Georgopoulos, C., Schrenzel, J., and Kelley, W.L. (2007). Isolation and characterization of biofilm formation-defective mutants of Staphylococcus aureus. Infect Immun 75, 1079–1088.

Wang, C.Y., Shi, J.D., Yang, P., Kumar, P.G., Li, Q.Z., Run, Q.G., Su, Y.C., Scott, H.S., Kao, K.J., and She, J.X. (2003). Molecular cloning and characterization of a novel gene family of four ancient conserved domain proteins (ACDP). Gene 306, 37–44.

Webb, B., and Sali, A. (2016). Comparative Protein Structure Modeling Using MODELLER. Curr Protoc Bioinformatics 54, 5 6 1–5 6 37.

Winter, G., Waterman, D.G., Parkhurst, J.M., Brewster, A.S., Gildea, R.J., Gerstel, M., Fuentes-Montero, L., Vollmar, M., Michels-Clark, T., Young, I.D., et al. (2018). DIALS: implementation and evaluation of a new integration package. Acta Crystallogr D Struct Biol 74, 85–97.

Wu, E.L., Cheng, X., Jo, S., Rui, H., Song, K.C., Davila-Contreras, E.M., Qi, Y., Lee, J., Monje-Galvan, V., Venable, R.M., et al. (2014). CHARMM-GUI Membrane Builder toward realistic biological membrane simulations. J Comput Chem 35, 1997–2004.

Yamazaki, D., Funato, Y., Miura, J., Sato, S., Toyosawa, S., Furutani, K., Kurachi, Y., Omori, Y., Furukawa, T., Tsuda, T., et al. (2013). Basolateral Mg2+ extrusion via CNNM4 mediates transcellular Mg2+ transport across epithelia: a mouse model. PLoS Genet 9, e1003983.

Yamazaki, D., Hasegawa, A., Funato, Y., Tran, H.N., Mori, M.X., Mori, Y., Sato, T., and Miki, H. (2019). Cnnm4 deficiency suppresses Ca(2+) signaling and promotes cell proliferation in the colon epithelia. Oncogene.

Yamazaki, D., Miyata, H., Funato, Y., Fujihara, Y., Ikawa, M., and Miki, H. (2016). The Mg2+ transporter CNNM4 regulates sperm Ca2+ homeostasis and is essential for reproduction. J Cell Sci 129, 1940–1949.

Yang, M., Jensen, L.T., Gardner, A.J., and Culotta, V.C. (2005). Manganese toxicity and Saccharomyces cerevisiae Mam3p, a member of the ACDP (ancient conserved domain protein) family. Biochem J 386, 479–487.

Zheng, H., Chordia, M.D., Cooper, D.R., Chruszcz, M., Muller, P., Sheldrick, G.M., and Minor, W. (2014). Validation of metal-binding sites in macromolecular structures with the CheckMyMetal web server. Nat Protoc 9, 156–170.

