## Supplementary Table S1 and Figures S1-S9 for "Crystal structure of a bacterial CNNM magnesium transporter"

### Supplemental Table and Figures

**Table S1.** Statistics of data collection and refinement

| | $MtCNNM_{CBS}$ + $Mg^{2+}$-ATP | $MtCNNM\Delta C_{\Delta loop}$ + $Mg^{2+}$-ATP | MtCNNMΔC R235L |
|---|---|---|---|
| **Data collection** | | | |
| X-ray source | ALS 5.0.2 | CLS 08ID-1 | ALS 5.0.2 |
| Wavelength (Å) | 1.00003 | 0.97996 | 1.03319 |
| Space group | $P4_122$ | $P2_12_12_1$ | C2 |
| Cell dimensions | | | |
| *a*, *b*, *c* (Å) | 52.10, 52.10, 112.28 | 61.05, 118.68, 177.31 | 139.37, 124.44, 85.27 |
| *α, β, γ* (°) | 90.0, 90.0, 90.0 | 90.0, 90.0, 90.0 | 90.0, 92.3, 90.0 |
| Resolution (Å) | 50.00–2.20 (2.24–2.20)[1] | 50.00–3.25 (3.31–3.25)[1] | 85.21-4.50 (5.03-4.50)[1] |
| Redundancy | 21.5 (11.2) | 11.9 (9.1) | 6 (5.7) |
| Completeness (%) | 99.0 (89.7) | 99.2 (96.9) | 99.9 (99.9) |
| $I/\sigma I$ | 40.6 (2.0) | 22.0 (1.0) | 4.8 (2.3) |
| $CC_{1/2}$ | 0.993 (0.780) | 0.993 (0.430) | 0.998 (0.852) |
| **Refinement** | | | |
| Resolution (Å) | 35.00–2.20 | 49.31–3.25 | 85.21–4.50 |
| No. of reflections | 7746 | 14835 | 8657 |
| $R_{work}/R_{free}$ | 0.213/0.243 | 0.242/0.279 | 0.334/0.345 |
| No. of atoms | | | |
| Protein | 929 | 4786 | 4145 |
| Ligands | 33 | 300 | 14 |
| Water | 51 | NA | NA |
| *B*-factors | | | |
| Protein | 38.6 | 24.9 | 174.62 |
| Ligands | 27.5 | 41.3 | 149.68 |
| Water | 39.7 | NA | NA |
| RMSDs | | | |
| Bond lengths (Å) | 0.002 | 0.002 | 0.001 |
| Bond angles (°) | 0.43 | 0.54 | 0.36 |
| Ramachandran plots | | | |
| Favored (%) | 97.5 | 96.2 | 94.6 |
| Allowed (%) | 2.5 | 3.8 | 4.7 |
| Disallowed (%) | 0.0 | 0.0 | 0.7 |
| **PDB code** | 7LJ6 | 7LJ7 | 7LJ8 |

[1]Highest resolution shell is shown in parentheses

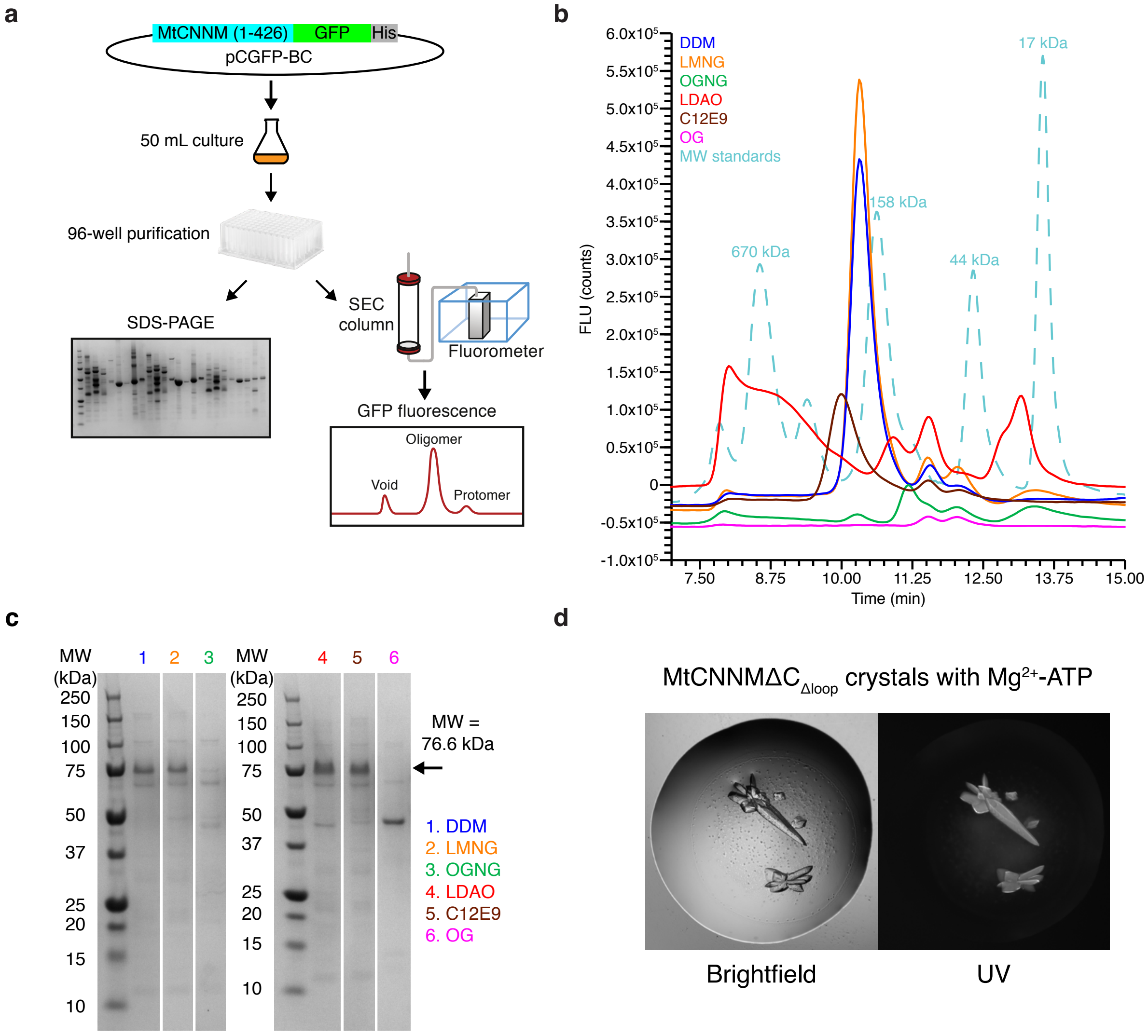


**Figure S1. Detergent screening and crystallization of MtCNNM. a**, Schematic of high-throughput screening process. **b**, Size-exclusion chromatography (SEC) profile of GFP-MtCNNM purified in different detergents. The dashed line shows molecular weight (MW) standards. **c**, SDS-PAGE analysis of GFP-MtCNNM purified in 6 detergents. **d**, MtCNNMΔC$_{\Delta loop}$ crystals in complex with $Mg^{2+}$-ATP taken with brightfield (*left*) and UV (*right*) camera.

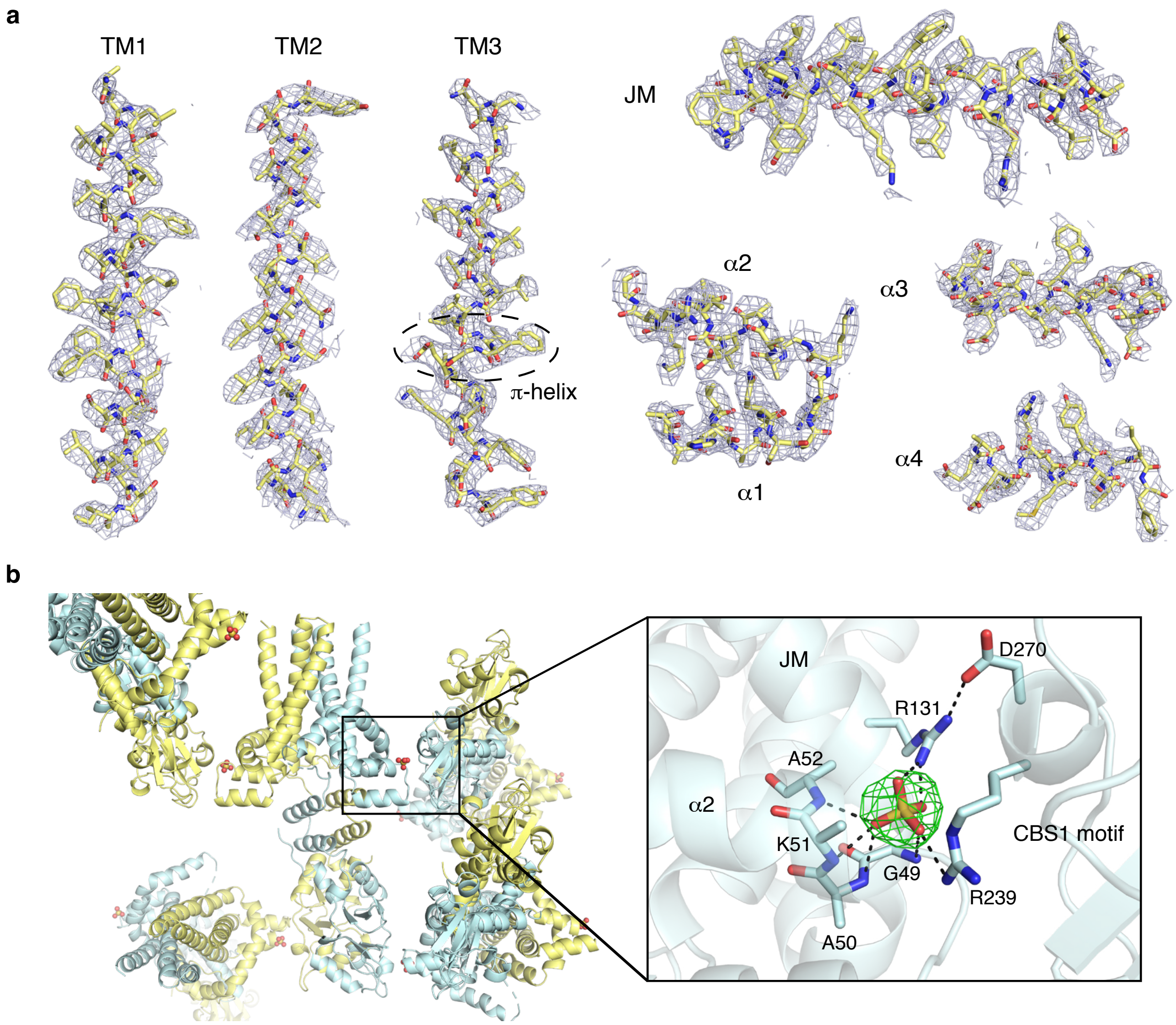


**Figure S2. Representative electron density. a**, Representative $2F_o$-$F_c$ map for TMD and AHB, contoured at 1.0 σ. **b**, Representative $F_o$-$F_c$ omit map for a sulfate ion between crystal packing surface, contoured at 5.0 σ.

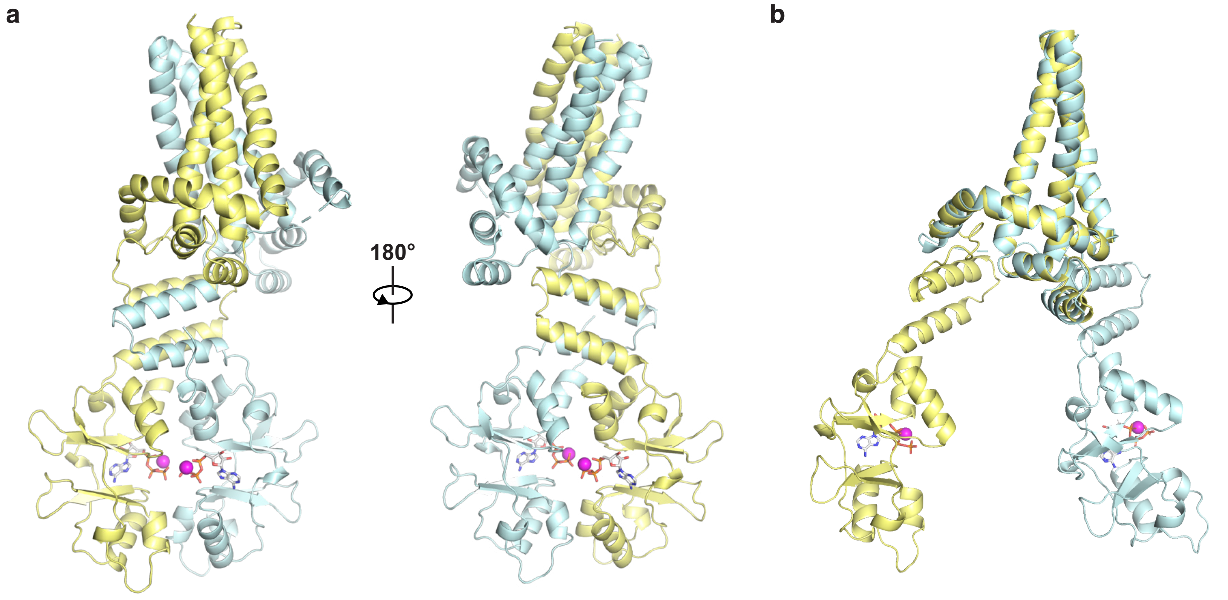


**Figure S3. Asymmetry between TMD and cytosolic domains in the protein crystal. a**, Front and rear view of the MtCNNMΔC homodimer showing asymmetry between TMD and cytosolic domains. **b**, Overlay of the TMD of the two protomers.

```
                      TM1                  α1              α2
Archaea     MVVIDLLIVEVVLFIAALLFSGFFSSSEVALISITRAKVHALQSQGRK----GAKALDTLKRST-DAI  63
Bacteria    --METSTIISLIIFILLIALTTVFVGSEFALVKIRATRIEQLADEGNK----PAKIVKKMIANL-DYY  61
Plant       SQVLKVLREQGLILAVLLGLSAFFSMAETSITTLWPWKVRELAEKEP-----ENGVFRMLRSDV-TRF  208
Yeast       VGDEASVTTYYIISIILVLLGGVFAGLTLGLMGQDEVYLKVISTSGSNSEKKLAKRVLDLISRGKHWV  122
Roundworm   KEYFLPLPLQIACIGFLLCLSALFSGLTLGLMSLTPQELELVIKSGAIKEQKCAAKILPVRKKG-NLL  261
Fruit fly   FEPLIPVWLAIIIIVTCLGFSALFSGLNLGLMSMDRTELKILRNTGTEKEKKYASKIAPVRDQG-NYL  246
Zebrafish   KKFLLPFWLQVIFIAMLLCLSGMFSGLNLGLMALDPMELRIVQNCGTEKEKHYAKAIEPVRSQG-NYL  258
Human       KKFLLPFWLQVIFISLLLCLSGMFSGLNLGLMALDPMELRIVQNCGTEKEKNYAKRIEPVRRQG-NYL  315
                             : :  .*     .:       :. :              .  :

                    TM2                                TM3
Archaea     QITTLIGSTIANVAVASLATAIGITLY--------------GNLGIAVGLVVAAVLVLVFGEIGPKMY  117
Bacteria    LSACQLGITVTSLGLGWLGEPTFEKLLHPIFEAINLPTALTTTISFAVSFIIVTYLHVVLGELAPKSI  129
Plant       LTTILIGTTVVNIAATALVTKAATAIF--------------GEAGVSAATGVMTVAILLLTEITPKSV  262
Yeast       LVTLLLSNVITNETLPIVLDRCL--------------------GGGWQAVVSSTILIVIFGEIIPQSV  170
Roundworm   LCSLLLGNVIVNSAISILMGEL---------------------TTGIYALIGSTMGIVIFGEILPQSI  308
Fruit fly   LCSILLGNVLVNSTFTILLDGL---------------------TSGLFAVIFSTLAIVLFGEITPQAV  293
Zebrafish   LCSLLLGNVLVNTTLTILLDDIA--------------------GSGLIAVVVSTIGIVIFGEIVPQAI  306
Human       LCSLLLGNVLVNTTLTILLDDIA--------------------GSGLVAVVVSTIGIVIFGEIVPQAI  363
              :  :. .:..     :                              .    :   :::.*: *:

                    JM                                      α3
Archaea     ASRYTEELALRVSRPILFFSKLLYPVLWVTDRIEQQF---AFR-PGVTEPVVTEEEIKEWIDVGEEEG  181
Bacteria    AIQHTEKLALVYARPLFYFGNIMKPLIWLMNGSARVIIRMFGVNPDAQTDAMSEEEIKIIINNSYNGG  197
Plant       AVHNAQEVARIVVRPVAWLSLILYPVGRVVTYLSMGILKILGL-KGRSEPYVTEDELKLMLRGAELSG  329
Yeast       CVKYGLQVGAFFCPFVLVLMYLMYPVAYPIATLLDYM---LGEDH---GTMYKKSGLKTLVTLHRTMG  232
Roundworm   CVKKGLEVGAHTISITQLFIFLTFPIAWPVSKLLDCL---LGDEY---Q-AYDRKRLMELIKMSITDN  369
Fruit fly   CSRHGLAIGAKTILVTKTVMAITAPLSYPVSRILDKL---LGEEI---GNVYNRERLKELVRVTNDVN  355
Zebrafish   CSRHGLAVGANTIFLTKFFMILTFPASYPVSKLLDHV---LGQEI---GTVYNREKLLEMLRVTDPYN  368
Human       CSRHGLAVGANTIFLTKFFMMMTFPASYPVSKLLDCV---LGQEI---GTVYNREKLLEMLRVTDPYN  425
            . :    :.          .   :   *          .               .. :   :

                α4           αA1  αA2                α5          β1
Archaea     TI--EEEERDMLYSVLRFGDTTAREVMTPRVDVVMIEDTATLES-ALAIFNETGFSRIPVYHE-RIDN  245
Bacteria    EI--NQTELAYMQNIFSFDERHAKDIMVPRTQMITLNEPFNVDE-LLETIKEHQFTRYPITDDGDKDH  262
Plant       AI--EEEEQDMIENVLEIKDTHVREVMTPLVDVVAIDGSGSLVD-FHNFWVTHQYSRVPVFEQ-RIDN  393
Yeast       VERLTKDEVTIISAVLDLKAKRVEEIMTPIENVFTMSADTILDDKTVEKIFNSGFSRIPIFLPNEPNN  300
Roundworm   GQ--VSNELKIAVGAMEIADKVVKDVMTKIEDVFMLPDTTVLNAKTVMEIVKMGYTRIPVYQYGDKNN  435
Fruit fly   DL--DKNEVNIISGALELRKKTVADVMTHINDAFMLSLDALLDFETVSEIMNSGYSRIPVYD-GDRKN  420
Zebrafish   DL--VKEELNIIQGALELRTKTVEDVMTPLRDCFMISGDAILDFATMSEIMESGYTRIPVYE-GERCH  433
Human       DL--VKEELNIIQGALELRTKTVEDVMTPLRDCFMITGEAILDFNTMSEIMESGYTRIPVFE-GERSN  490
               . *             : :    . ::*.    :  . :     :               ::*  *:              :

            β2   α6                 αB       β3         α7           β4
Archaea     IVGLLNVKDVFSAVFRQQ--TSATIRDL-MYEPYFIPESKKIDELLKELQVKK-QHMAVVLDE-----  304
Bacteria    IKGFINVKEFLTEYASGKTIKIAN---YIH-ELPMISETTRISDALIRMQREH-VHMSLIIDE-----  320
Plant       IVGIAYAMDLLDYVPKGKLLESTTVVDMAHKPAFFVPDSMSVWNLLREFRIRK-VHMAVVLNE-----  455
Yeast       FIGMLLVRVLISYDPDDCL-PISHFP---LATLPETSPNTSCLNILNYFQEGK-AHMCVVSKE-----  358
Roundworm   VTDMLFVKDLALLDPDDNF-TVKTVCGYHKHPVKFVMNDTPLPNLLEAFKKGE-GHLAMVKRLINTDD  501
Fruit fly   IVTLLYIKDLAFVDTDDNT-PLKTLCEFYQNPVHFVFEDYTLDIMFNQFKEGTIGHIAFVHRV-NNEG  486
Zebrafish   IVDLLFVKDLAFVDPDDCT-PLKTITKFYSHPLHFVFNDTKLDAMLEEFKKGK-SHLAIVQRV-NNEG  498
Human       IVDLLFVKDLAFVDPDDCT-PLKTITKFYNHPLHFVFNDTKLDAMLEEFKKGK-SHLAIVQRV-NNEG  555
            .  :      .                                          :   ::      *:..:

                     β5   α8
Archaea     ----YGSFAGIVTVEDMLEELV  322
Bacteria    ----YGGTAGILTMEDILEEIV  338
Plant       ----YGGTIGIVTLEDVVEEIV  473
Yeast       -PGSSHGAIGVLTLEDVIEELI  379
Roundworm   KHDPSYVLVGVVTLEDIVEEIL  523
Fruit fly   DGDPFYETVGLVTLEDVIEELI  508
Zebrafish   EGDPFYEVLGIVTLEDVIEEII  520
Human       EGDPFYEVLGIVTLEDVIEEII  577
                     *::*:**::**::
```

polar residues in acidic cavity

polar residues in outward-facing conformation

ATP-binding site residues

**Figure S4. Sequence alignment of CNNM orthologs from eight representative species.** The listed CNNM orthologs and their UniProt accession numbers are: *Methanoculleus thermophilus* (A0A1G8XA46), *Staphylococcus aureus* (A0A0H3JL60), *Arabidopsis thaliana* (Q84R21), *Saccharomyces cerevisiae* (Q12296), *Caenorhabditis elegans* (A3QM97), *Drosophila melanogaster* (A0A0B7P9G0), *Danio rerio* (A2ATX7), and *Homo sapiens* (Q9H8M5). Secondary structure corresponds to the crystal structure of MtCNNMΔC. Highlighted residues: polar residues in acidic cavity (*magenta*), polar residues in outward-facing conformation (*green*), and ATP-binding site residues (*orange*).

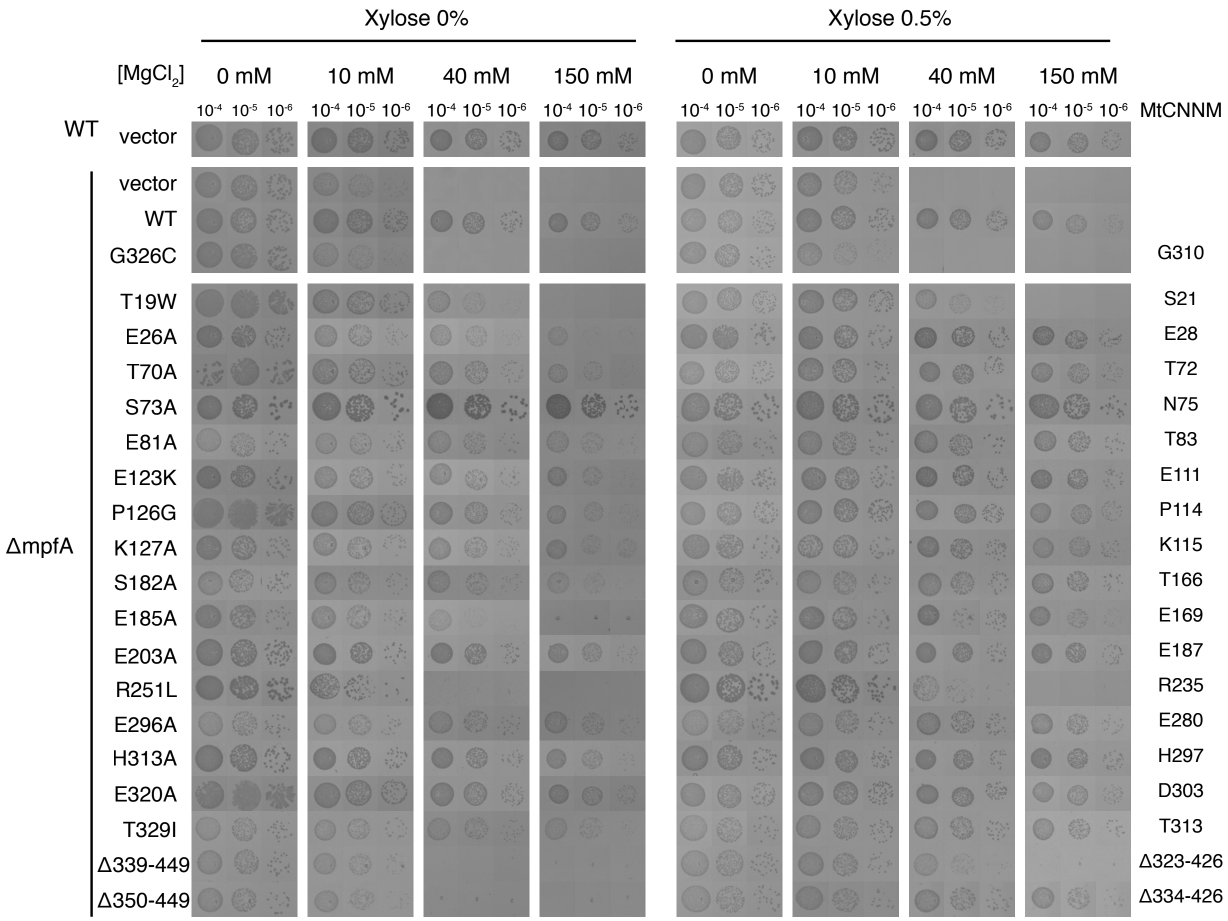


**Figure S5. In vivo complementation assay of MpfA with various mutants in high magnesium condition.** The *S. aureus* ΔmpfA strain was transformed with various plasmids. Serially diluted overnight cultures were spotted on plates with varying $MgCl_2$ and xylose concentration. Inactive mutants are unable to grow under high magnesium condition.

a

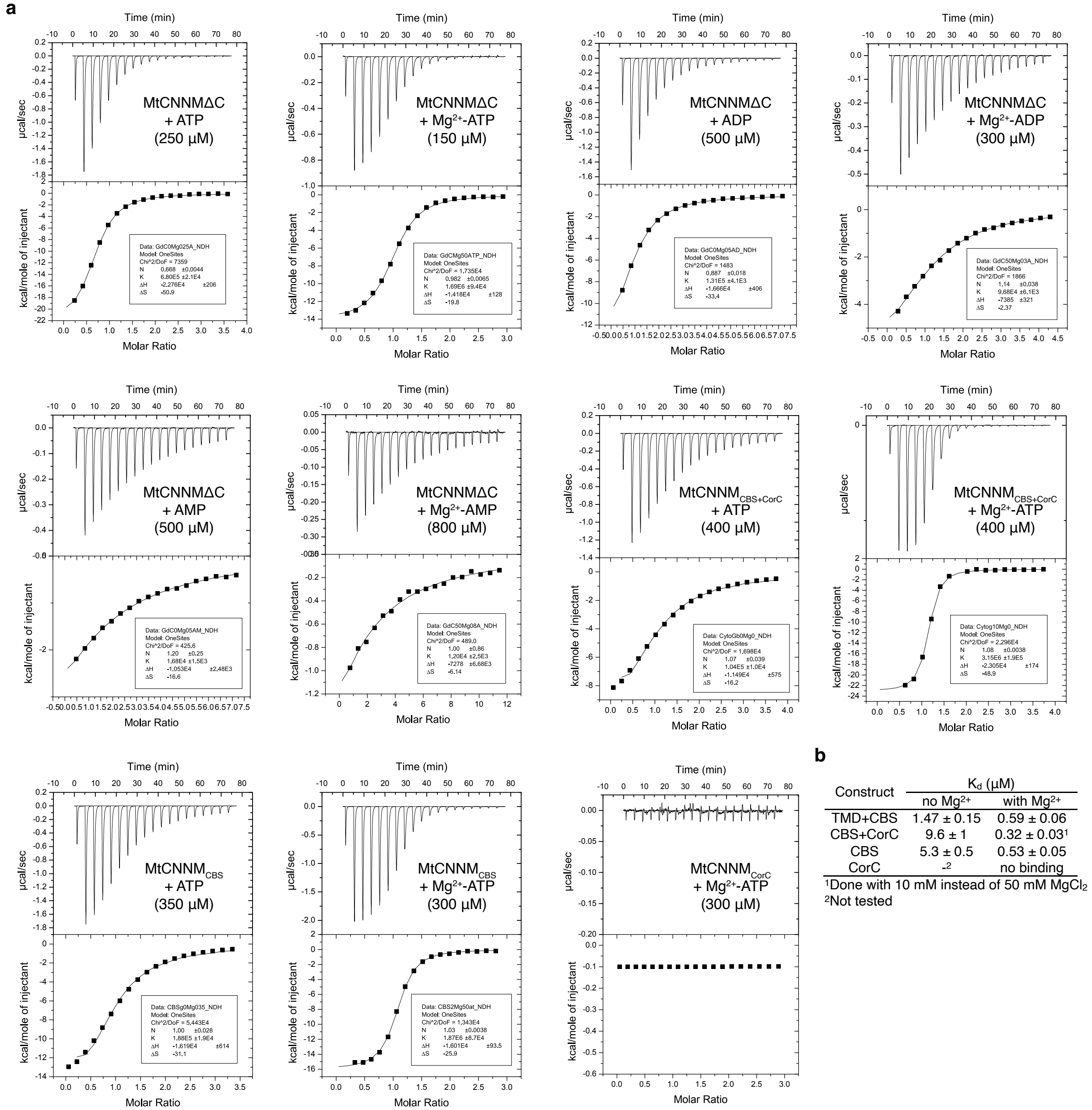


b

| Construct | $K_d$ (µM) no $Mg^{2+}$ | $K_d$ (µM) with $Mg^{2+}$ |
|---|---|---|
| TMD+CBS | 1.47 ± 0.15 | 0.59 ± 0.06 |
| CBS+CorC | 9.6 ± 1 | 0.32 ± 0.03[1] |
| CBS | 5.3 ± 0.5 | 0.53 ± 0.05 |
| CorC | -[2] | no binding |

[1]Done with 10 mM instead of 50 mM $MgCl_2$
[2]Not tested

**Figure S6. Raw ITC thermograms. a**, ITC thermograms showing various MtCNNM constructs binding to adenosine nucleotides (concentrations indicated in parentheses) in absence or presence of 50 mM $MgCl_2$. **b**, Table summarizing the affinities of different MtCNNM constructs binding to ATP in presence or absence of 50 mM $MgCl_2$.

**a**

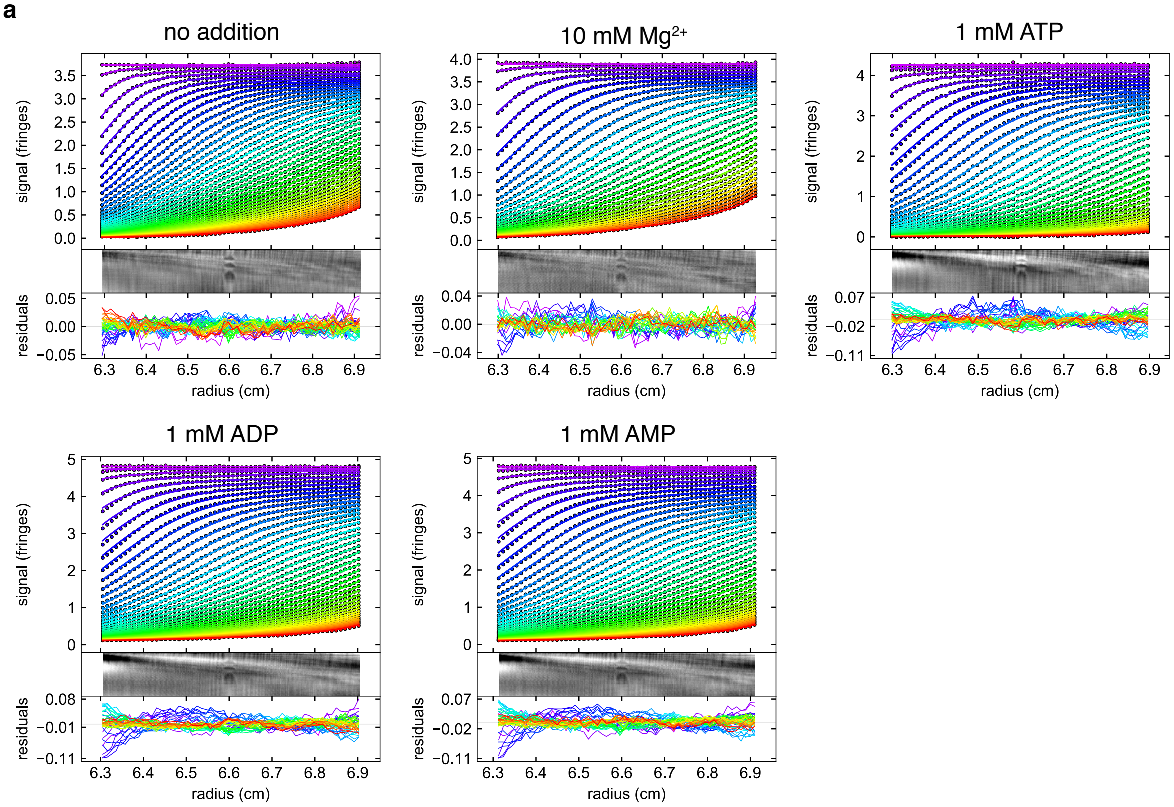


**b**

| | Sedimentation coefficient (S) | | Estimated MW (kDa) | | RMSD | Theoretical MW (kDa) | |
|---|---|---|---|---|---|---|---|
| Condition | Peak 1 | Peak 2 | Peak 1 | Peak 2 | | Monomer | Dimer |
| no addition | 1.84 | - | 16.2 | | 0.0049 | | |
| 10 mM $Mg^{2+}$ | 1.78 | - | 16.0 | - | 0.0042 | | |
| 1 mM ATP | - | 2.49 | - | 32.0 | 0.0045 | 14.5 | 29.0 |
| 1 mM ADP | - | 2.40 | - | 30.4 | 0.0044 | | |
| 1 mM AMP | - | 2.44 | - | 30.9 | 0.0050 | | |

**Figure S7. Summary of SV-AUC results. a**, Sedimentation velocity AUC profiles of $MtCNNM_{CBS}$ in presence of various ligands. Interference of the sample are plotted against the radial position in the cell. One in every 75 scans is plotted. **b**, Summary of experimental sedimentation coefficients and estimated molecular weights.

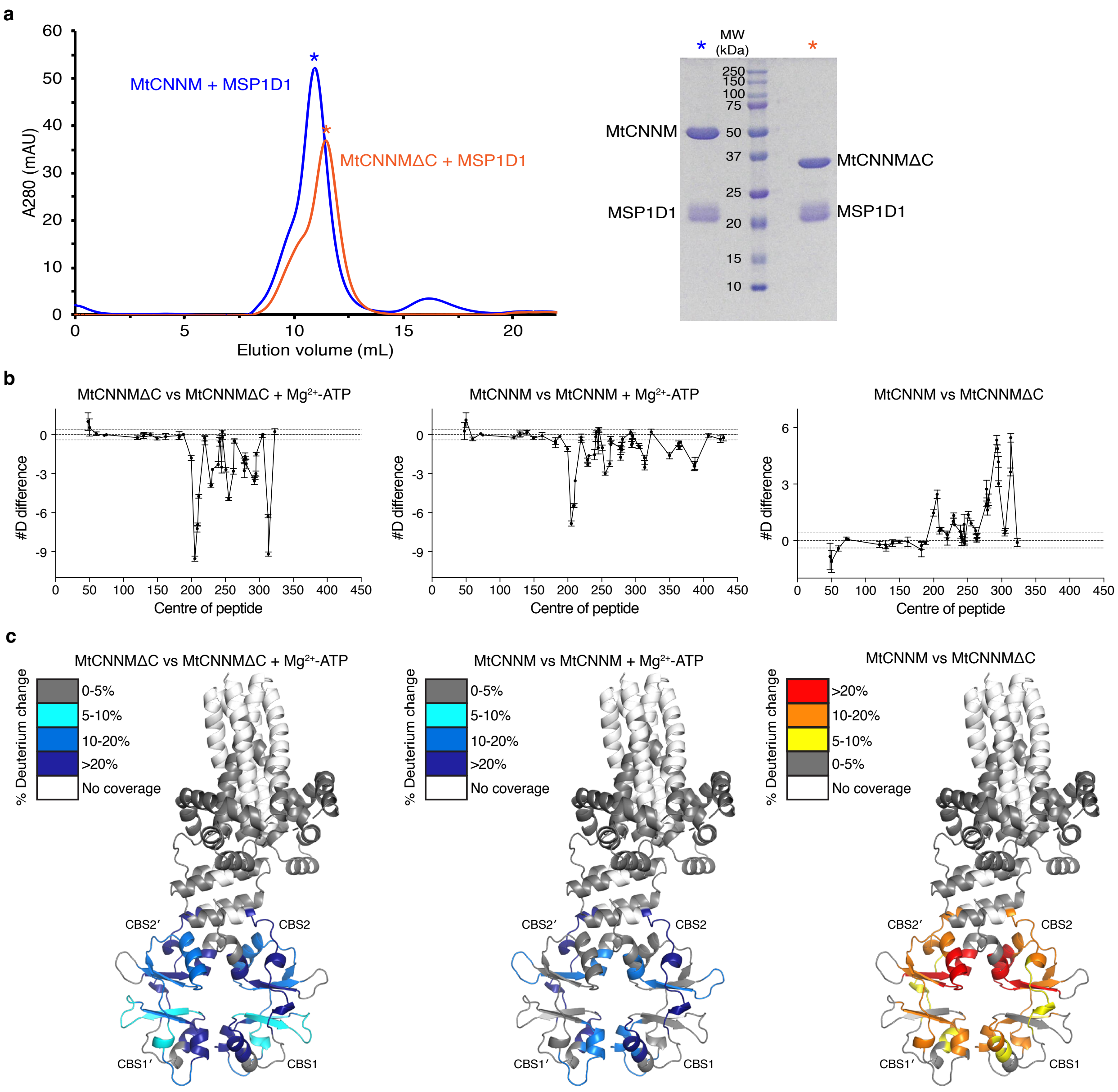


**Figure S8. Reconstitution of MtCNNM into MSP1D1 nanodiscs and HDX-MS analyses. a**, SEC profile and SDS-PAGE analysis of MtCNNM and MtCNNMΔC reconstituted in MSP1D1 nanodiscs. **b**, HDX-MS analysis of three sets of experiments. The sum of the # of deuterons protected from across all timepoints is shown. Each point represents a single peptide, with them being graphed on the x-axis according to its central residue. Error bars represent standard deviation (n = 3). **c**, Mapping of deuteration onto the structure of MtCNNMΔC. Regions that showed significant decreases or increases in exchange (defined as >5%, 0.3 kDa, and a Student's *t*-test $p<0.01$) are colored in blue or red respectively.

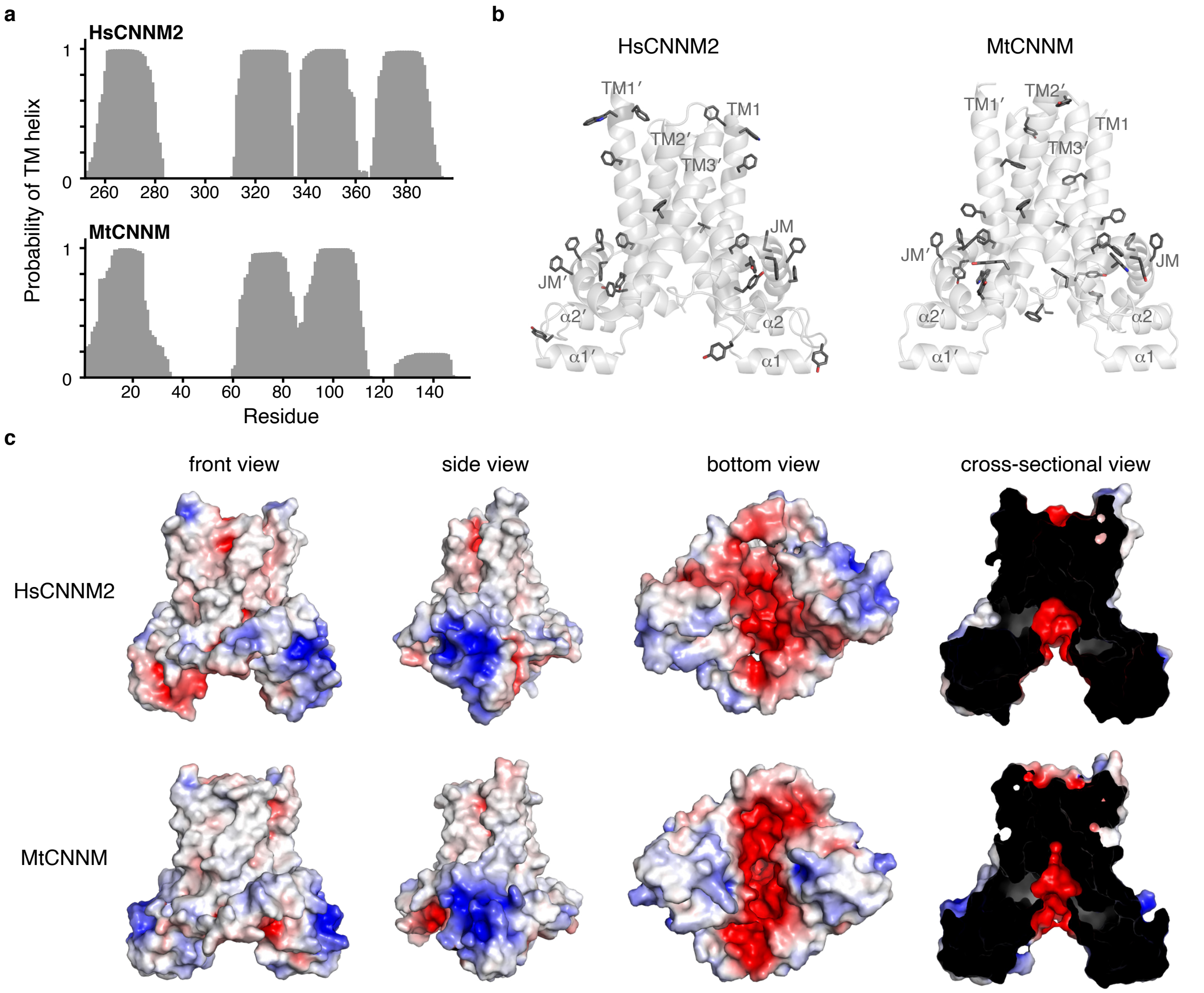


**Figure S9. Analysis of HsCNNM2 model structure. a,** Prediction of transmembrane helices in human and archaeal CNNM with TMHMM server. HsCNNM2 is predicted to have four TM helices. **b**, Clustering of aromatic residues at the phospholipid-solvent interface and JM helix. **c,** Comparison of the electrostatic surfaces of human and archaeal TMD (±5 kT $e^{-1}$).

**Supplemental Video 1: Unbiased MD simulations of MtCNNMΔC showing movements of cytosolic domains.**

**Supplemental Video 2: Targeted MD simulations of MtCNNM TMD from inward-facing to outward-facing conformations.**

**Supplemental Data 1: This file contains the source data for the HDX-MS analyses shown in Fig. S8.**
